# Ipsilateral stimulus encoding in primary and secondary somatosensory cortex of awake mice

**DOI:** 10.1101/2021.07.16.452704

**Authors:** Aurélie Pala, Garrett B. Stanley

## Abstract

Lateralization is a hallmark of somatosensory processing in the mammalian brain. However, in addition to their contralateral representation, unilateral tactile stimuli also modulate neuronal activity in somatosensory cortices of the ipsilateral hemisphere. The cellular organization and functional role of these ipsilateral stimulus responses in awake somatosensory cortices, especially regarding stimulus coding, are unknown. Here, we targeted silicon probe recordings to the vibrissa region of primary (S1) and secondary (S2) somatosensory cortex of awake head-fixed mice of either sex while delivering ipsilateral and contralateral whisker stimuli. Ipsilateral stimuli drove larger and more reliable responses in S2 than in S1, and activated a larger fraction of stimulus-responsive neurons. Ipsilateral stimulus-responsive neurons were rare in layer 4 of S1, but were located in equal proportion across all layers in S2. Linear classifier analyses further revealed that decoding of the ipsilateral stimulus was more accurate in S2 than S1, while S1 decoded contralateral stimuli most accurately. These results reveal substantial encoding of ipsilateral stimuli in S1 and especially S2, consistent with the hypothesis that higher cortical areas may integrate tactile inputs across larger portions of space, spanning both sides of the body.

**S1ignificance Statement:** Tactile information obtained by one side of the body is represented in the activity of neurons of the opposite brain hemisphere. However unilateral tactile stimulation also modulates neuronal activity in the other, or ipsilateral, brain hemisphere. This ipsilateral activity may play an important role in the representation and processing of tactile information, in particular when the sense of touch involves both sides of the body. Our work in the whisker system of awake mice reveals that neocortical ipsilateral activity, in particular that of deep layer excitatory neurons of secondary somatosensory cortex (S2), contains information about the presence and the velocity of unilateral tactile stimuli, which supports a key role for S2 in integrating tactile information across both body sides.

## Introduction

Most studies of somatosensation concentrate on a single cerebral hemisphere and examine the neocortical representations of tactile signals arising from the opposite, or contralateral, side of the body. However, across species, ipsilateral tactile stimuli have also been shown to evoke changes in population activity of primary (S1) and secondary (S2) somatosensory cortex (Pidoux and Verley 1979, Tommerdahl et al. 2005, Hlushchuk and Hari 2006, Lipton et al. 2006, Ferezou et al. 2007, Eickhoff et al. 2008, Plomp et al. 2017, Song et al. 2018), mainly mediated by corticocortical projections via the corpus callosum (Pidoux and Verley 1979, Picard et al. 1990, Fabri et al. 1999). Yet, surprisingly little is known about the cellular-level specificity of ipsilateral stimulus-evoked activity in S1 and S2, and about its potential role in the neocortical encoding of tactile information during awake somatosensation.

Previous studies of ipsilateral activity in somatosensory cortices have focused on putative excitatory neurons, revealing sensory responses distinct from contralateral ones. Ipsilateral stimulation of the hand in macaque monkeys and of the whiskers in rodents primarily elicited increased spiking in subsets of putative excitatory neurons in S1 (in area 2 in monkeys) (Iwamura et al. 1994, Iwamura et al. 2001, Shuler et al. 2001, Wiest et al. 2005) and S2 (Carvell and Simons 1986, Burton et al. 1998, Iwamura et al. 2001, Taoka et al. 2016). These sensory responses were typically smaller, sparser, and exhibited longer onset latency than sensory responses evoked by contralateral stimuli. In comparison, GABAergic neuron responses to ipsilateral tactile stimuli have not been investigated (but see Palmer et al. 2012), even though fast-spiking (FS) GABAergic neurons have been shown to receive interhemispheric callosal inputs in vitro (Petreanu et al. 2007, Karayannis et al. 2007, Rock and Apicella 2015) and in vivo (Cisse et al. 2003, Cisse et al. 2007) in multiple neocortical areas.

Separately, anatomical studies have revealed differences in the density of callosal axon terminals as a function of the neocortical lamina they innervate in both S1 and S2 (Wise 1975, Wise and Jones 1976, Akers and Killackey 1978, Sloper and Powell 1979, Petreanu et al. 2007). Yet, whether sensory responses evoked by ipsilateral stimuli exhibit laminar-specific organization potentially suggestive of an intracortical subnetwork dedicated to ipsilateral tactile information processing is completely unexplored.

In addition to its cellular organization, a major unknown pertaining to S1 and S2 activity relates to its role in ipsilateral stimulus coding. Whether changes in population spiking enable the decoding of ipsilateral tactile stimuli, and whether this differs in S1 and S2, is totally unknown. Previous studies on the encoding of contralateral whisker stimuli have revealed that the spike rate of single neurons and of populations of neurons in the vibrissa region of S1 and of S2 support the prediction of the stimulus occurrence, or its detection (Wang et al. 2010, Adibi and Arabzadeh 2011, Kwon et al. 2016). S1 spikes are also known to encode contralateral stimulus properties, for instance enabling the discrimination between whisker deflections of different amplitudes (Adibi and Arabzadeh 2011), different velocities (Wang et al. 2010), and of different temporal profiles (Arabzadeh et al. 2006, McGuire et al. 2016). To what extent ipsilateral stimuli can be detected and discriminated from S1 and S2 activity, given that they elicit weaker, sparser and delayed changes in spiking, is uncertain.

Here, we reveal substantial representation of ipsilateral stimuli in the neural activity of awake S1 and especially S2. We first show that ipsilateral stimuli evoke larger and more reliable sensory responses in a larger fraction of putative excitatory neurons (Regular-Spiking, RS) and FS inhibitory neurons, with less laminar specificity in S2 compared to S1. Then, we reveal that increased and decreased RS spiking in both S1 and S2 enables ipsilateral stimulus detection and stimulus velocity discrimination, with S2 spiking showing higher ipsilateral stimulus detectability and discriminability. These results suggest that S2 may be key in integrating both contralateral and ipsilateral tactile signals.

## Materials and Methods

### Animals, headpost implantation, and habituation to head restraint

Thirteen 8-26 week old C57BL/6J male mice, one 11 week old male and one 12 week old female Scnn1a-Tg3-Cre (Madisen et al. 2010) mouse crossed with CAG-LSL- ChR2(H134R)-EYFP mice (LSL-ChR2) (Madisen et al. 2010) were used in accordance with protocols approved by the Georgia Institute of Technology Institutional Animal Care and Use Committee and in agreement with guidelines established by the National Institutes of Health. Mice were housed in groups of two individuals (minimum) under a reversed light-dark cycle. Mice were implanted with a custom-made headpost and a recording chamber under 1-1.5% isoflurane anesthesia. After minimum 3 days of recovery, mice were gradually habituated to head fixation, paw restraint, and whisker stimulation for 3-6 days before proceeding to electrophysiological recordings.

### Identification and verification of recording location

Primary (S1) and secondary (S2) somatosensory cortex recording locations were functionally identified via intrinsic signal optical imaging (ISOI) performed through a thinned skull under 1-1.25% isoflurane anesthesia (Yamashita et al. 2013, Masino et al.1993). S1 and S2 recordings were mainly targeted to areas corresponding to the B1 and B2 whiskers (18 S1 recordings in total: B1 whisker: 10 recordings, B2 whisker: 2 recordings, C1 whisker: 3 recordings, C2 whisker: 1 recording, D2 whisker: 2 recordings / 20 S2 recordings in total: B1 whisker: 14 recordings, C1 whisker: 3 recordings, C2 whisker: 3 recordings). We pooled the data obtained from recordings targeted to areas corresponding to different whiskers, since these did not differ in the fraction of ipsilateral stimulus-responsive RS neurons, nor in the RS neuron change in spiking evoked by ipsilateral stimuli in either S1 and S2. Additionally, we verified the precise location and the insertion angle and depth of the silicon probes by imaging the fluorescent probe tracks in fixed brain slices stained to highlight layer 4 across S1 and S2. In brief, after the last recording, mice were transcardially perfused with 1x PBS (137 mM NaCl, 2.7 mM KCl, and 10 mM phosphate buffer, VWR), followed by 4% paraformaldehyde. The brains were extracted and post-fixed for a maximum of 2 hours in the 4% paraformaldehyde solution before being sectioned in 100 um thick coronal slices on a vibratome. The brain slices were stained for cytochrome oxidase activity to highlight the location of S1 barrel cortex (Wong-Riley and Welt 1980) and of S2 layer 4, before being further incubated with DAPI (2 µM in PBS) for 15 minutes, mounted on slides with Fluoromount, and imaged using a confocal microscope.

### Silicon probe recordings

Mice were anesthetized (1-1.5% isoflurane anesthesia), and a small craniotomy was made above the left hemisphere at the exact location previously determined by ISOI (see above) leaving the dura intact. The craniotomy was then covered with silicone elastomer (Kwik-Cast, WPI) and mice were returned to their home cage for at least 2 hours to recover from anesthesia. In a subset of mice, recordings were conducted in the same craniotomy across two consecutive days. Mice were placed on the recording setup, the silicone elastomer removed, and a 32-channel laminar silicon probe (A1x32-5mm-25- 177-A32, 25 µm inter-channel spacing, Neuronexus) was slowly inserted through the dura using a micromanipulator (Luigs & Neumann) to a target depth of 1000-1100 µm. The probe insertion angle was 35° from the vertical for S1 recordings, and 55° for S2 recordings. All silicon probes were electrochemically plated with a poly(3,4- ethylenedioxythiophene) (PEDOT) polymer (Wilks et al. 2009, Ludwig et al. 2011) using a NanoZ device (White Matter LLC) to reach 1 kHz impedance values between 0.2 and 0.5 MΩ. Silicon probes were then coated with DiI (0.2 mg/ml in ethanol) (Invitrogen) to be able to visualize their fluorescent track in fixed tissue after the termination of the recordings (see above). Once the silicon probe was lowered to its target depth, a drop of agarose gel (2% in Ringer solution) (Sigma) was applied on top of the craniotomy to minimize movements and prevent drying of the recording site, followed by a drop of mineral oil to prevent drying of the agarose. Data collection started after a minimum of 30 minutes to allow for relaxation of the brain tissue. Continuous signals were filtered (1st- order high-pass at 0.3 Hz and 3rd-order low-pass at 7.5 kHz) and digitized at 30 kHz using a 128-channel Cerebus system (Blackrock Microsystems).

### Whisker stimulation

All but three whiskers from distinct rows on each side of the face were trimmed at their base. The left and right whiskers corresponding to the recorded region of S1 or S2 were threaded into narrow 1.5 cm long extension tubes glued to high-precision galvanometer-operated stimulators (Cambridge Technologies) under the control of custom routines written in MATLAB and Simulink Real-Time (Mathworks) with 1 ms temporal resolution. The other four whiskers (two on each side of the face) were imaged to identify epochs of whisker stillness (see below). The extension tubes were positioned approximately 5 mm away from the face and aligned with the whisker natural resting orientations. Whiskers were deflected in the caudo-rostral direction following a sawtooth-shaped spatiotemporal profile (Wang et al. 2010). The rise and decay times of the sawtooth waveform were 8 ms, and deflection velocity was calculated as the average velocity across the whole waveform duration (16 ms). Left and right whiskers were randomly stimulated with a minimum of 2 s between consecutive stimuli.

### Whisker movement videography and identification of epochs of “Whisker stillness”

For most recordings, videography was acquired for two non-trimmed whiskers on each side of the face (see above) at 200 Hz with a resolution of 14.4 pixels/mm (EoSens CL MC1362, Mikrotron), while in a subset of recordings, whisker videography was acquired at 25 Hz with a resolution of 6.8 pixels/mm (HQCAM), under infrared illumination. The identification of epochs of whisker stillness and of whisking was done using custom routines written in MATLAB (MathWorks). In brief, the movie pixel grayscale values were first inverted such that the whiskers appeared white on a darker background. Then, one region of interest (ROI) was manually delineated on each side of the face, and the absolute across-frames variation of the normalized sum of the pixel values within each ROI was calculated and then summed across the two ROIs. The obtained time series was then smoothed and individual time points with values lower than a fixed threshold were labeled as “Whisker stillness”. “Whisker stillness” epochs shorter than 25 ms (5 frames at 200 Hz) were removed from the “Whisker stillness” category.

### Layer 4 depth estimation in Scnn1a-Tg3-Cre x LSL-ChR2 mice

To confirm the accuracy of our functional laminar estimation (see below), we performed two S1 and four S2 recordings in two Scnn1a-Tg3-Cre mice crossed with CAG-LSL-ChR2(H134R)-EYFP mice, which express the Channelrhodopsin-2 protein (ChR2) in layer 4 excitatory neurons of S1 and S2 (Madisen et al. 2010, Pluta et al. 2015, Minamisawa et al. 2018). ChR2 excitation was achieved with a 470 nm LED (Thorlabs) coupled to a 400 µm diameter optic fiber (Thorlabs) placed immediately above the craniotomy. The pattern of light stimulation was a train of square light pulses of 3 ms duration and 19.1 mW/mm^2^ intensity delivered with a minimum of 1 s inter-pulse interval. The center of L4 was assigned to the silicon probe channel fulfilling the largest number of the following four criteria: 1) time of peak of the light-evoked local field potential (LFP) response within 2 % of the fastest peak time across all 32 channels, 2) peak amplitude of the light-evoked LFP response within 95 % of the largest peak amplitude across all 32 channels, 3) sink peak times of the current source density (CSD) analysis of the light-evoked LFP response within twice the fastest CSD sink peak time across all 32 channels, and 4) sink onset in the CSD within twice the fastest CSD sink onset time across all 32 channels (Sofroniew et al. 2015). Details regarding the LFP and CSD stimulus-evoked response calculations are described below as they are similar for the responses evoked by light and whisker stimuli. The identity of the silicon probe channel assigned to the center of L4 was then compared to that obtained using our sensory response-based method (see below).

### Electrophysiology data analysis

All electrophysiology data analyses were conducted in MATLAB (MathWorks).

#### Spike sorting and identification of single-unit clusters

Individual recording sweeps were band-passed filtered (forward and reverse, 4^th^ order Butterworth filter, cutoff frequencies of 500 Hz and 14.25 kHz) and concatenated before proceeding to automated spike sorting using Kilosort2 (Pachitariu et al. 2016) and manual curation of the spike clusters using Phy (Rossant and Harris 2013).

Spike clusters were assigned to the channel with the largest trough-to-peak amplitude (Voltage Trough-to-Peak, VTP), measured on the cluster average spike waveform. Spike clusters were considered as single-unit if they met the following six criteria: 1) more than 500 individual spikes in the cluster, 2) signal-to-noise (SNR) ratio of the average spike waveform larger than 5. SNR was defined as the ratio between the trough-to-peak amplitude and the mean standard deviation across the entire duration (3 ms) of the waveform, 3) coefficient of variation (CV) across the whole recording duration of the VTP averaged over 120 s windows smaller than 0.2, 4) CV across the whole recording duration of the spiking rate averaged over 120 s windows smaller than 1, 5) fraction of inter-spike intervals shorter than 2 ms, or refractory period violations, smaller than 1% (Fee et al. 1996, Hill et al. 2011), 6) cluster isolation distance larger than 55. Isolation distance was calculated as the Mahalanobis distance between the *n*^th^ closest non-cluster spike waveform to the cluster spike waveforms, with *n* being the number of spikes in the cluster (Harris et al. 2001). Each cluster and non-cluster spike waveform were described using the first three principal components across all channels. The single-unit clusters included in subsequent analyses contained on average 31108 ± 40622 spikes (mean ± SD), had a SNR of 8.0 ± 2.7, a VTP CV of 0.069 ± 0.041, a spike rate CV of 0.39 ± 0.18, a fraction of refractory period violations of 0.17 % ± 0.20 %, and an isolation distance of 93 ± 81. On average, 27 single-unit clusters were isolated per recording.

Regular Spiking (RS) putative excitatory neurons were distinguished from fast-spiking (FS) putative inhibitory neurons on the basis of the time elapsed from trough to peak (TtoP) of the average cluster waveform. Clusters with a TtoP value smaller than 0.4 ms were identified as FS neurons, while clusters with TtoP values larger than 0.5 ms were labeled as RS neurons (Bartho et al. 2004, Sofroniew et al. 2015). Clusters with TtoP values in the 0.4-0.5 ms range were not included in the analyses. Using such metric and thresholds, 76% of the single-unit clusters were classified as RS neurons (263 neurons in S1, 359 neurons in S2) and 21% as FS neurons (74 neurons in S1, 98 neurons in S2).

#### Layer 4 and individual neurons depth estimation

To estimate the depth of L4 in S1 and S2 recordings we considered the LFP, CSD and multi-unit (MUA) responses evoked by contralateral whisker stimuli (Sederberg et al. 2019). The average LFP response was obtained by down-sampling to 3 kHz and low-pass filtering the raw signal (forward and reverse, 200 Hz cutoff frequency). The one-dimensional CSD was calculated from the second spatial derivative of the average LFP response (Freeman and Nicholson 1975) with sinks having negative values and sources positive values. For display, the CSD profiles were interpolated along the depth axis. The average MUA response was obtained by high-pass filtering (3^rd^ order Butterworth filter, 800 Hz cutoff frequency), rectifying and smoothing the raw signal. The center of L4 was assigned to the silicon probe channel fulfilling the largest number of the following four criteria (Haslinger et al. 2006, Higley and Contreras 2007, Plomp et al. 2014): 1) LFP response peak time within 2 % of the fastest LFP peak response time across all 32 channels, 2) LFP response peak amplitude within 95 % of the largest LFP peak response amplitude across all 32 channels, 3) sink onset in the CSD within 2 % of the fastest CSD sink onset time across all 32 channels, 4) MUA response onset time within 2 % of the fastest MUA onset response time across all 32 channels. The thickness of layer 4 was estimated as 200 µm in S1, equivalent to 8 channels on the silicon probe, and 175 µm in S2, equivalent to 7 channels, according to our own measurements in fixed tissue sections and consistent with prior studies (Hooks et al. 2011). Individual neuron depth equaled the depth of the channel to which they were assigned (see above), leading to 5 %, 12 %, and 83 % of S1 RS neurons, and 18 %, 17 %, and 65 % of S2 RS neurons recorded in L2/3, L4, and L5/6 respectively, matching previously reported proportions in rodent neocortex (Naka et al. 2019, Horvath et al. 2021).

#### Sensory response quantification

Mean sensory responses were obtained by averaging individual sensory responses evoked by whisker stimuli occurring during epochs of whisker stillness. Responses were included in the average only if the 80 ms prior and the 160 ms after the onset of the whisker stimulus were assigned to the “Whisker stillness” category (see above). Across recordings, 57 % ± 12 % (mean ± SD, range: 25 % – 78 %) of the stimuli occurred during epochs of whisker stillness, resulting in 95 ± 37 stimulus trials (mean ± SD, range: 23 – 182 trials) used to calculate the mean response evoked by either ipsilateral or contralateral stimuli.

The magnitude of sensory responses was calculated by subtracting the mean spike rate calculated over a 500 ms window immediately prior to stimulus onset – the baseline firing rate – from the mean spike rate calculated over a 50 ms window starting at stimulus onset. The z-scored magnitude was obtained by dividing the mean response magnitude by the standard deviation of the baseline firing rate across stimulus trials. Sensory response variability was estimated by calculating the coefficient of variation (CV) of the response magnitude across stimulus trials, that is by dividing the standard deviation of the response magnitude by the absolute value of the mean response magnitude. The onset latency of positive and negative sensory responses was defined as the earliest time-point post stimulus onset for which the baseline-subtracted cumulative PSTH was above or below a 95% bootstrapped confidence interval on the cumulative baseline values (Wiest et al. 2005). Only onset latencies shorter than 50 ms were included in the population analyses.

All single-neuron and population PSTHs had a bin size of 1 ms. Population PSTHs had their overall pre-stimulus baseline spike rate calculated over a 500 ms window immediately prior to stimulus onset subtracted from every bin value before being smoothed by convolution with a Gaussian function with 2-ms standard deviation.

#### Identification of stimulus-responsive neurons

A neuron was considered stimulus-responsive if it met two out of the three following criteria: 1) for a PSTH with 10 ms bin size, at minimum 2 bins within the first 50 ms post-stimulus onset with a value above, or 4 bins with a value below, a 95% confidence threshold on the pre-stimulus spike rate obtained by bootstrapping, 2) a bootstrapped 95% confidence interval on the mean response magnitude (see above) that did not include 0 spikes/sec, 3) different spike count distributions for a post- and a pre-stimulus epoch of 50 ms duration at a significance level of 0.05 assessed by a one-tailed Wilcoxon rank-sum test. Further, criteria 1) and 3) determined whether sensory responses were positive or negative. For evaluating stimulus-responsiveness, responses to all whisker stimuli, irrespective of the presence or absence of whisker movements at the time of stimulus delivery, were included in the analysis.

#### Spike count correlation

Spike count correlation (rSC) between pairs of RS neurons was computed as the Pearson correlation coefficient between the number of spikes occurring in a 50 ms window starting immediately post stimulus onset for repeated presentations of the stimulus. Only trials where the stimulus occurred during epochs of whisker stillness were used in the analysis, and no other trial selection criteria were used. Spike count correlation was computed for all possible pairs of RS neurons, irrespective of their stimulus responsiveness. To compare rSC values in S1 and S2, rSCs were converted to *z*-scores using the Fisher transformation.

### Linear Discriminant Analysis (LDA) classifiers

To assess the detectability of the 1000 °/s whisker stimuli from S1 and S2 RS neuron activity, spiking data from 8 S1 and 8 S2 recordings with a minimum of 10 simultaneously recorded RS neurons each and at least 75 stimulus trials occurring during epochs of whisker stillness (see above) were used. The classifier input population was either made of all simultaneously recorded RS neurons in a given recording (*within-recording* classifier) (S1: 22 ± 4 RS/rec, median ± MAD, range: 14 – 35 RS/rec, S2: 27 ± 9 RS/rec, range: 14 – 40 RS/rec), or of a selection of RS neurons randomly sampled across all S1 or all S2 recordings (*across-recordings* classifier) (selection pool size of 184 RS neurons for S1 and 205 RS neurons for S2). For each neuron, stimulus trials were partitioned into 10 folds. For the *across-recordings* classifiers, 90 trials were sampled with replacement from 9 of the folds to create a training set, while 10 trials were sampled with replacement from the remaining fold to create a testing test. For each “Stim” trial, the number of spikes occurring in a 50 ms window located immediately post stimulus onset was used as input to the classifier, while the number of spikes occurring during a similar duration window located immediately before stimulus onset was used as input to the classifier for the “No Stim” trials. This led to a total of 126 trials (63 “Stim” and 63 “No Stim”) in the training set and 14 trials (7 “Stim” and 7 “No Stim”) in the testing set for the *within-recording* classifiers, and 180 training trials (90 “Stim” and 90 “No Stim”) and 20 test trials (10 “Stim” and 10 “No Stim”) for the *across-recordings* classifiers. The LDA classifier was trained on the trials of the training set using a full covariance matrix for the *within-recording* classifiers and a diagonal covariance matrix for the *across-recordings* classifiers, while classification accuracy was evaluated on the testing set. The procedure was repeated until all folds were used to generate the testing set, and mean classification accuracy was calculated by averaging classification accuracy values obtained for each of the 10 distinct testing/training trial partitions. For the *across-recordings* classifiers, the neuron selection process followed by classifier training and testing according to a 10-fold cross-validation scheme was repeated 100 times, and the median classification accuracy with a bootstrapped estimate of the median standard deviation was reported. Chance level classification accuracies were obtained by randomly shuffling the labels (“Stim” or “No Stim”) of the trials of the training set.

To assess the detectability of the 1000 °/s whisker stimuli from the activity of S1 and S2 RS neurons located in different neocortical layers, the same procedure as described above for the *across-recordings* classifiers was followed. A random selection of 10 RS neurons was used as the classifier input population to account for the selection pool size of each layer (S1: 11 L2/3 RS neurons, 24 L4 RS neurons, 149 L5/6 RS neurons, S2: 35 L2/3 RS neurons, 34 L4 RS neurons, 135 L5/6 RS neurons).

To investigate the contribution of stimulus-responsive neurons with positive and negative sensory response magnitude to the detectability of 1000 °/s whisker stimuli, we repeated the same procedure as described above for the *across-recordings* classifiers, while varying the initial pool from which 24 RS neurons were selected. We chose a classifier input population size of 24 neurons, as it reflected the average number of simultaneously recorded RS neurons across the 16 S1 and S2 recordings included in the classification analysis. The S1 pool sizes were 61 S1 RS neurons for stimulus-responsive neurons (R), 25 RS neurons for stimulus-responsive neurons with positive response magnitude (R>0), 36 RS neurons for stimulus-responsive neurons with negative response magnitude (R>0), and 123 RS neurons for non-stimulus-responsive neurons (no R). The S2 pool sizes were 74 RS neurons (R), 33 RS neurons (R>0), 41 RS neurons (R<0), and 107 RS neurons (no R).

To assess the detectability of the 200 °/s whisker stimuli from S1 and S2 RS neuron activity, 6 S1 and 6 S2 recordings with a minimum of 10 simultaneously recorded RS neurons and at least 75 stimulus trials occurring during epochs of whisker stillness (see above) were used to generate a pool of 169 S1 neurons and 144 S2 neurons out of which the LDA classifiers were built and their performance evaluated as described above for the *across-recordings* classifier.

To assess the discriminability of 200 °/s vs 1000 °/s whisker stimuli from S1 and S2 RS neuron activity, 3 S1 and 3 S2 recordings with a minimum of 10 simultaneously recorded RS neurons and at least seventy-five 200 °/s and seventy-five 1000 °/s stimulus trials occurring during epochs of whisker stillness (see above) were used to generate a pool of 83 S1 neurons and 77 S2 neurons out of which the LDA classifiers were built (*across-recordings* classifiers). The procedure to train and evaluate the classifiers was similar to that used for probing stimulus detectability, except that the inputs to the classifier were spike counts measured over a 50 ms window immediately post stimulus onset for both 200 °/s and 1000 °/s trials.

All classifier-based analyses were conducted in MATLAB (MathWorks).

### Experimental design and statistical analysis

We carried out non-parametric Wilcoxon rank-sum and Wilcoxon signed-rank tests to compare the median of two distributions of unpaired and paired samples respectively, except for comparing spike count correlation distributions, where we used a *t*-test. Chi-squared tests were used to assess differences between proportions of neurons. LDA classifier performances were compared using Wilcoxon rank-sum tests. When more than two comparisons were performed between more than two groups, Bonferroni correction was used to adjust the significance levels of the statistical tests. A minimum of 1000 bootstrap samples were generated to produce confidence intervals and to estimate the standard deviation of the median in all analyses involving *across-recordings* classifiers. All statistical analyses were conducted in MATLAB (MathWorks).

### Data availability

Source data and code to reproduce the analyses and figures can be downloaded from the Zenodo repository (10.5281/zenodo.5899625).

## Results

### S2 neurons exhibit more frequent, larger and less variable sensory responses to ipsilateral stimuli

We performed laminar silicon probe recordings in vibrissa S1 and S2 of the left hemisphere of awake, head-restrained, mice. We simultaneously measured the spiking activity of populations of individual putative excitatory neurons (Regular-Spiking, RS) (Figure 1) and fast-spiking inhibitory neurons (FS) (Figure 2) in response to 1000 °/s punctate deflections of a single somatotopically-aligned ipsilateral whisker. For comparison, we applied the same single-whisker stimuli to the somatotopically-aligned contralateral whisker. To avoid any modulation of stimulus-evoked changes in spike rate by whisker movements (Fanselow and Nicolelis 1999), we focused all our analyses on stimuli delivered when the whiskers were immobile as determined by high-speed videography (Figure 1A).

**Figure 1.**
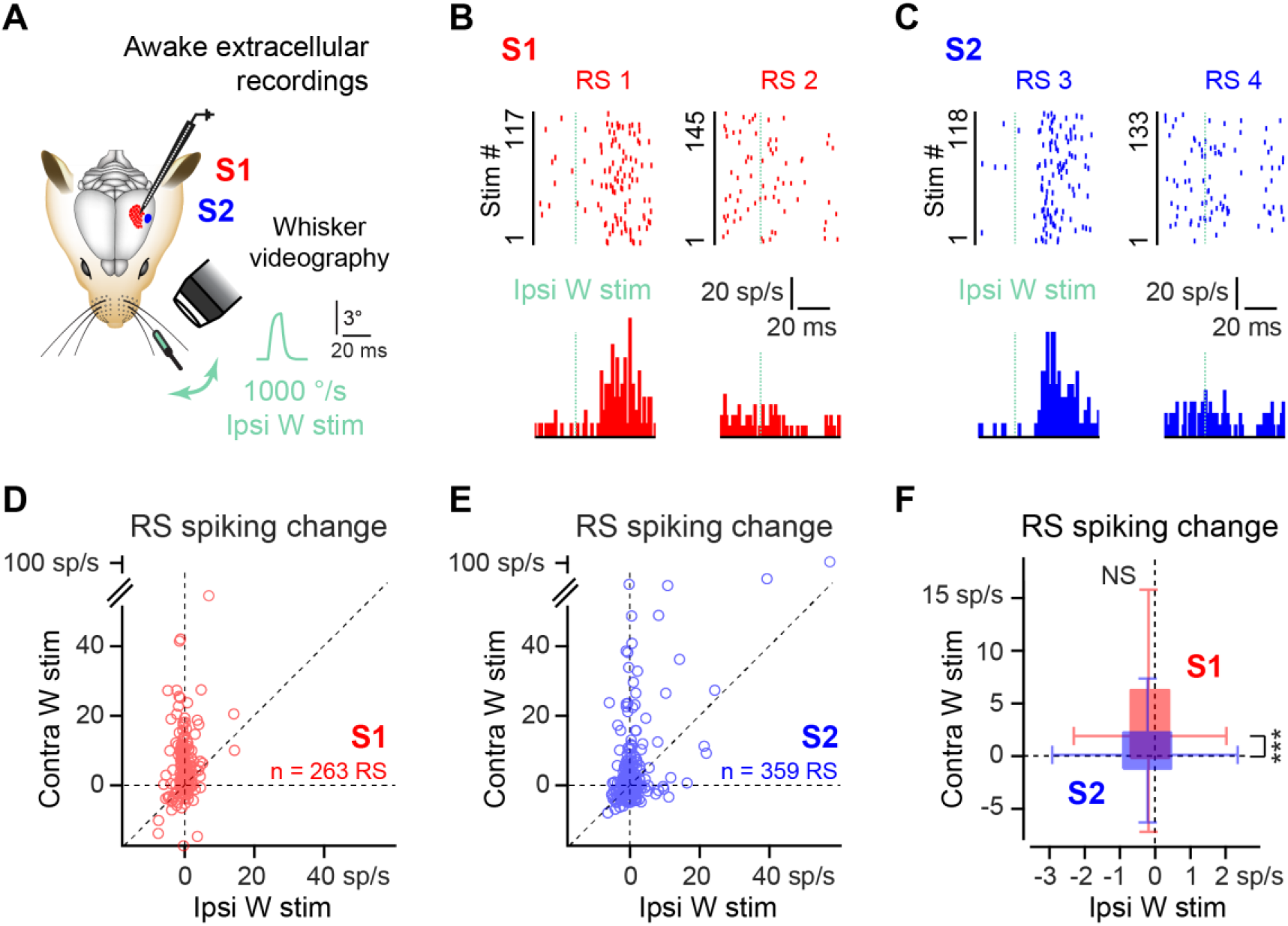
Change in S1 and S2 RS neuron spiking evoked by ipsilateral whisker stimuli. (A) Change in spiking activity evoked by 1000 °/s deflections of the somatotopically-aligned ipsilateral whisker is measured through laminar silicon probe recordings. High-speed videography is used to confirm the absence of whisker movements before and after stimulation. (**B**) Example spike raster plots and PSTHs for two S1 RS neurons with increased (left) and decreased (right) spiking in response to ipsilateral whisker stimulation. (**C**) Same as (B), but for two S2 RS neurons. (**D**) Mean spike rate change evoked in 263 S1 RS neurons by ipsilateral stimuli with corresponding contralateral stimulus-evoked spike rate change. (**E**) Same as (D) for 359 S2 RS neurons. (**F**) Ipsilateral stimuli elicit a decrease in spike rate of comparable amplitude in S1 and S2 RS neurons (S1: -0.19 ± 0.55 spikes/s (n=263), median ± MAD, p = 0.0013, two-sided sign test, S2: -0.22 ± 0.69 spikes/s (n=359), p = 5.23·10^- 5^, S1 vs S2: p = 0.68, two-sided Wilcoxon rank-sum test). Contralateral stimuli elicit a larger change in spike rate in S1 RS neurons compared to S2 RS neurons (S1: 1.48 ± 2.50 spikes/s (n=263), median ± MAD, p = 1.22·10^-10^, two-sided sign test, S2: 0.11 ± 1.61 spikes/s (n=359), p = 0.40, S1 vs S2: p = 1.02·10^-6^, two-sided Wilcoxon rank-sum test). NS p≥0.05, ** p<0.01, *** p<0.001.

**Figure 2.**
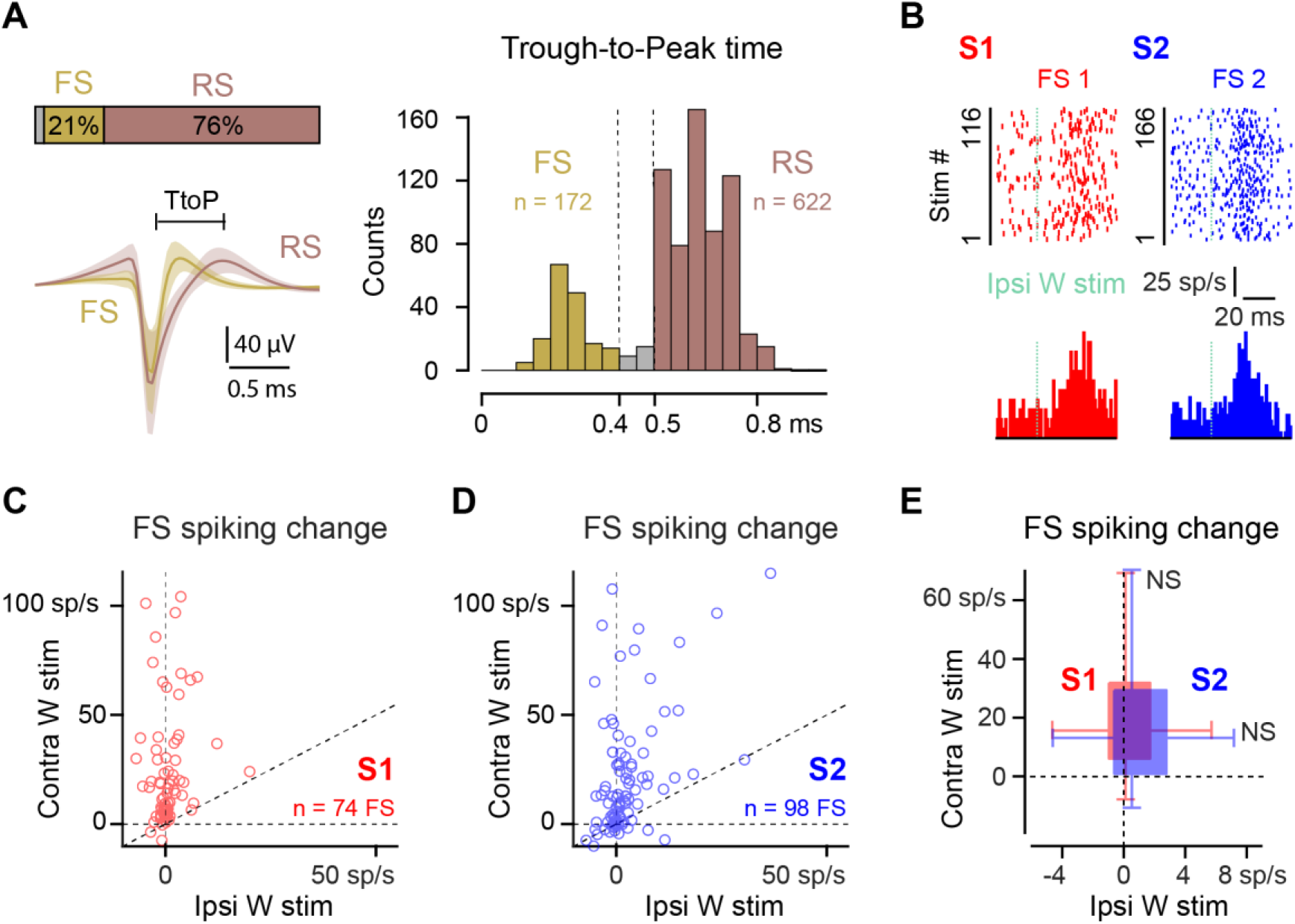
Change in S1 and S2 FS neuron spiking evoked by ipsilateral whisker stimuli. **(A)** FS neurons are identified by a spike waveform Trough-to-Peak (TtoP) time shorter than 0.4 ms. and RS neurons by a TtoP longer than 0.5 ms. (**B**) Example spike raster plots and PSTHs for one S1 (red) and one S2 (blue) FS neuron with increased spiking in response to ipsilateral whisker stimulation. (**C**) Mean spike rate change evoked in 74 S1 FS neurons by ipsilateral stimuli with corresponding contralateral stimulus-evoked spike rate change. (**D**) Same as (C) for 98 S2 FS neurons. (**E**) Ipsilateral stimuli elicit an increase in spike rate in S2 FS neurons (S1: 0.11 ± 1.29 spikes/s (n=74), median ± MAD, p = 0.48, two-sided sign test, S2: 0.61 ± 1.72 spikes/s (n=98), p = 0.032, S1 vs S2: p = 0.098, two-sided Wilcoxon rank-sum test). Contralateral stimuli elicit an increase in spike rate of similar amplitude in S1 and S2 FS neurons (S1: 14.84 ± 10.29 spikes/s (n=74), median ± MAD, p = 7.16·10^- 18^, two-sided sign test, S2: 12.49 ± 12.51 spikes/s (n=98), p = 2.87·10^-9^, S1 vs S2: p = 0.12, two-sided Wilcoxon rank-sum test). NS p≥0.05, * p<0.05, *** p<0.001.

First, we characterized the effect of stimulating the ipsilateral whisker on the spiking activity of RS neurons of S1 and S2. Ipsilateral stimuli drove both increases and decreases in RS neuron spiking relative to ongoing activity in S1 (Figure 1B) and S2 (Figure 1C), resulting in heterogenous effects across S1 and S2 RS neuron populations (Figure 1D, E). Overall, ipsilateral stimuli elicited a small but significant reduction in RS neuron spike rate in both S1 and S2 (S1: -0.19 ± 0.55 spikes/s (n=263), median ± MAD, p = 0.0013, two-sided sign test, S2: -0.22 ± 0.69 spikes/s (n=359), p = 5.23·10^-5^), with no difference in magnitude between the two regions (p = 0.68, two-sided Wilcoxon rank-sum test) (Figure 1F). In comparison, and as expected, deflections of the contralateral whisker at the same velocity led to a notable increase in RS neuron spike rate in S1 that was significantly larger than in S2 (Figure 1F).

Contrary to the observations in RS neurons, ipsilateral stimuli mainly induced increased spiking in individual FS neurons (Figure 2A, see Materials and Methods) of S1 and S2 (Figure 2B-D), resulting in an overall positive change in FS neuron spike rate, of comparable magnitude across S1 and S2 (Figure 2E). Contralateral stimuli also elicited an increase in spike rate in both S1 and S2 FS neurons, but of much larger magnitude than the change in spiking evoked by ipsilateral stimuli (Figure 2C-E). A marked difference between S1 and S2 was the larger proportion of ipsilateral stimulus-responsive RS and FS neurons in S2 compared to S1 (RS: S1: 31 % (82/263), S2: 39 % (140/359), p = 0.0443, chi-squared test, FS: S1: 36 % (27/74), S2: 64 % (63/98), p = 0.00030). This was opposite to contralateral stimulus-responsive RS and FS neurons, which were more numerous in S1 than S2 (RS: S1: 85 % (223/263), S2: 71 % (256/359), p = 7.90·10^-5^, chi- squared test, FS: S1: 99 % (73/74), S2: 89 % (87/98), p = 0.012) (Table 1). Interestingly, amongst the population of ipsilateral stimulus-responsive RS neurons we found an equal proportion of neurons with positive (R>0) and negative (R<0) response magnitudes (S1: R>0: 44 % (36/82), R<0: 56 % (46/82), p = 0.12, chi-squared test, S2: R>0: 46 % (65/140), R<0: 54 % (75/140), p=0.23), whereas contralateral stimuli drove mainly positive RS neuron responses; further, FS neuron responses to either ipsilateral or contralateral stimuli were predominantly positive (Table 1).

**Table 1:** Proportion of stimulus-responsive RS and FS neurons and response properties: 1000 °/s stimuli. Proportion of RS and FS neurons with a significant response to 1000 °/s ipsilateral or contralateral stimuli. Response magnitude (z-score), variability (coefficient of variation, CV), and onset latency are reported as median ± MAD.

|  |  | Ipsi-responsive RS |  | Contra-responsive RS |  |
| --- | --- | --- | --- | --- | --- |
|  |  | R > 0 | R < 0 | R > 0 | R < 0 |
| **S1** | Proportion of neurons | 31% (82/263) |  | 85% (223/263) |  |
|  |  | 44% (36/82) | 56% (46/82) | 83% (184/223) | 17% (39/223) |
|  | Z-scored magnitude | 0.51 ± 0.20 | -0.36 ± 0.13 | 1.62 ± 1.06 | -0.47 ± 0.20 |
|  | Variability (CV) | 5.86 ± 2.58 | 4.25 ± 1.69 | 2.57 ± 1.37 | 3.11 ± 1.23 |
|  | Onset latency (ms) | 22.0 ± 7.5 | 19.0 ± 11.0 | 8.7 ± 1.7 | 25.0 ± 7.0 |
| **S2** | Proportion of neurons | 39% (140/359) |  | 71% (256/359) |  |
|  |  | 46% (65/140) | 54% (75/140) | 68% (173/256) | 32% (83/256) |
|  | Z-scored magnitude | 1.06 ± 0.64 | -0.40 ± 0.15 | 1.30 ± 1.05 | -0.49 ± 0.16 |
|  | Variability (CV) | 4.14 ± 2.14 | 3.59 ± 1.35 | 3.25 ± 2.08 | 2.79 ± 0.84 |
|  | Onset latency (ms) | 16.8 ± 4.2 | 25.0 ± 13.0 | 11.0 ± 2.0 | 23.0 ± 8.5 |
|  |  | Ipsi-responsive FS |  | Contra-responsive FS |  |
|  |  | R > 0 | R < 0 | R > 0 | R < 0 |
| **S1** | Proportion of neurons | 36% (27/74) |  | 99% (73/74) |  |
|  |  | 74% (20/27) | 26% (7/27) | 100% (73/73) | 0% (0/73) |
| **S2** | Proportion of neurons | 64% (63/98) |  | 89% (87/98) |  |
|  |  | 78% (49/63) | 22% (14/63) | 90% (78/87) | 10% (9/87) |

We thus further examined ipsilateral responses across S1 and S2 by separating positive (Figure 3A) and negative (Figure 3E) stimulus-responsive RS neurons (Table 1).

**Figure 3.**
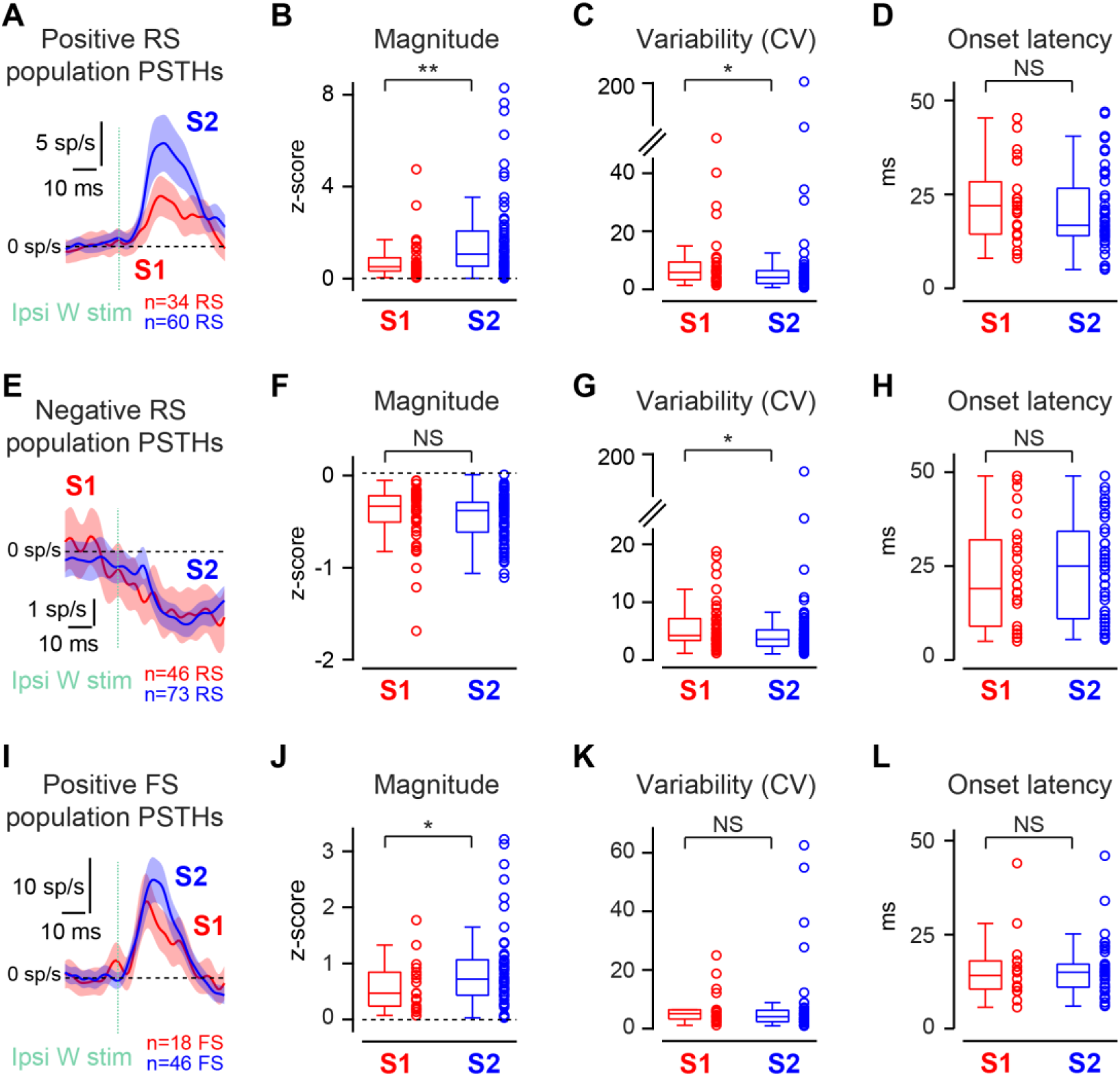
S2 neurons have larger and less variable responses to ipsilateral stimuli compared to S1 neurons. (**A**) Baseline-subtracted population PSTHs for S1 (red) and S2 (blue) RS neurons with positive responses to ipsilateral stimuli. Shaded area represents the SEM. (**B**) Larger magnitude of positive ipsilateral responses in S2 compared to S1 RS neurons (S1: 0.51 ± 0.20 (n=34), S2: 1.06 ± 0.64 (n=60), z-score, median ± MAD, p = 0.0025, one-sided Wilcoxon rank-sum test). (**C**) Smaller trial-to-trial variability of positive ipsilateral responses in S2 compared to S1 RS neurons. Variability is quantified as the coefficient of variation (CV) of the response magnitude (S1: 5.86 ± 2.58 (n=34), S2: 4.14 ± 2.14 (n=60), median ± MAD, p = 0.026, one-sided Wilcoxon rank-sum test). (**D**) Comparable onset latency for positive ipsilateral responses in S1 and S2 RS neurons (S1: 22.0 ± 7.5 ms (n=25), S2: 16.8 ± 4.2 ms (n=48), median ± MAD, p = 0.11, one-sided Wilcoxon rank-sum test). (**E**) Same as (A), but for RS neurons with negative responses to ipsilateral stimuli. (**F**) Magnitude of negative ipsilateral responses is comparable in S1 and S2 RS neurons (S1: -0.36 ± 0.13 (n=46), S2: -0.40 ± 0.15 (n=73), z-score, median ± MAD, p = 0.079, one-sided Wilcoxon rank-sum test). (**G**) Variability of negative ipsilateral responses is smaller in S2 compared to S1 RS neurons (S1: 4.25 ± 1.69 (n=46), S2: 3.59 ± 1.35 (n=73), median ± MAD, p = 0.014, one-sided Wilcoxon rank-sum test). (**H**) Comparable onset latency for negative ipsilateral responses in S1 and S2 RS neurons (S1: 19.0 ± 11.0 ms (n=26), S2: 25.0 ± 13.0 ms (n=49), median ± MAD, p = 0.31, one-sided Wilcoxon rank-sum test). (**I**) Same as (A), but for FS neurons with positive responses to ipsilateral stimuli. (**J**) Larger magnitude of positive ipsilateral responses in S2 compared to S1 FS neurons (S1: 0.47 ± 0.28 (n=18), S2: 0.72 ± 0.30 (n=46), z-score, median ± MAD, p = 0.048, one-sided Wilcoxon rank-sum test). (**K**) Comparable ipsilateral response variability in S1 and S2 FS neurons. (S1: 5.10 ± 1.59 (n=18), S2: 4.03 ± 1.67 (n=46), median ± MAD, p = 0.16, one-sided Wilcoxon rank-sum test). (**L**) Comparable onset latency for positive ipsilateral responses in S1 and S2 FS neurons (S1: 14.1 ± 3.9 ms (n=16), S2: 15.0 ± 4.0 ms (n=39), median ± MAD, p = 0.42, one-sided Wilcoxon rank-sum test). NS p≥0.05, * p<0.05, ** p<0.01.

Positive responses in RS neurons were larger in S2 than S1 (Figure 3B), and a similar but non-significant trend was observed for negative responses (Figure 3F). Regardless of response sign, response variability for RS neurons was smaller in S2 than S1 (quantified by the coefficient of variation (CV) of the response magnitude across repeated whisker stimulations) (Figure 3C, G) and onset latency was comparable in S1 and S2 (Figure 3D, H).

Notably, positive ipsilateral responses in S1 and S2 RS neurons had longer onset latencies than positive responses to contralateral stimuli (S1: Ipsi: 22.0 ± 7.5 ms (n=25), Contra: 8.7 ± 1.7 ms (n=160), median ± MAD, p = 1.73·10^-8^, two-sided Wilcoxon rank-sum test, S2: Ipsi: 16.8 ± 4.2 ms (n=48), Contra: 11.0 ± 2.0 ms (n=143), p = 6.56·10^-8^) (Table 1), similarly to what was previously reported for putative excitatory neurons of layer 5 in S1 (Shuler et al. 2001, Wiest et al. 2005). FS neurons, which mainly displayed positive responses to ipsilateral stimuli (Figure 3I) (Table 1), showed significantly larger response magnitude (Figure 3J) accompanied by smaller yet non-significant response variability (Figure 3K) in S2 compared to S1, and comparable onset response latency in both areas (Figure 3L). Taken together, these results show that ipsilateral whisker stimuli elicited larger and more reliable sensory responses in a larger fraction of RS and FS neurons in S2 compared to S1, therefore suggesting a more robust representation of the ipsilateral tactile inputs in S2 than in S1. We next examined layer-specific distributions and response profiles for ipsilateral stimulus-responsive RS and FS neurons in S1 and S2. We found a smaller proportion of ipsilateral stimulus-responsive neurons in L4 of S1 as compared to L2/3 and to L5/6 (Figure 4A, B), while in S2 stimulus-responsive neurons were found in equal proportions across all layers (Figure 4C, D), consistent with prior anatomical studies on the laminar location of callosal axon terminals (Wise 1975, Wise and Jones 1976, Akers and Killackey 1978, Petreanu et al. 2007). However, positive and negative sensory response magnitude (Figure 4E), variability (Figure 4F), and onset latency (Figure 4G), in RS and FS neurons were comparable across laminae in both S1 and S2. This suggests that the representation of the ipsilateral tactile inputs is widely distributed across neurons of all layers in S2, while in S1, ipsilateral responses spare L4, the main thalamocortical recipient layer, which may rather be dedicated to representing and processing contralateral tactile information.

**Figure 4.**
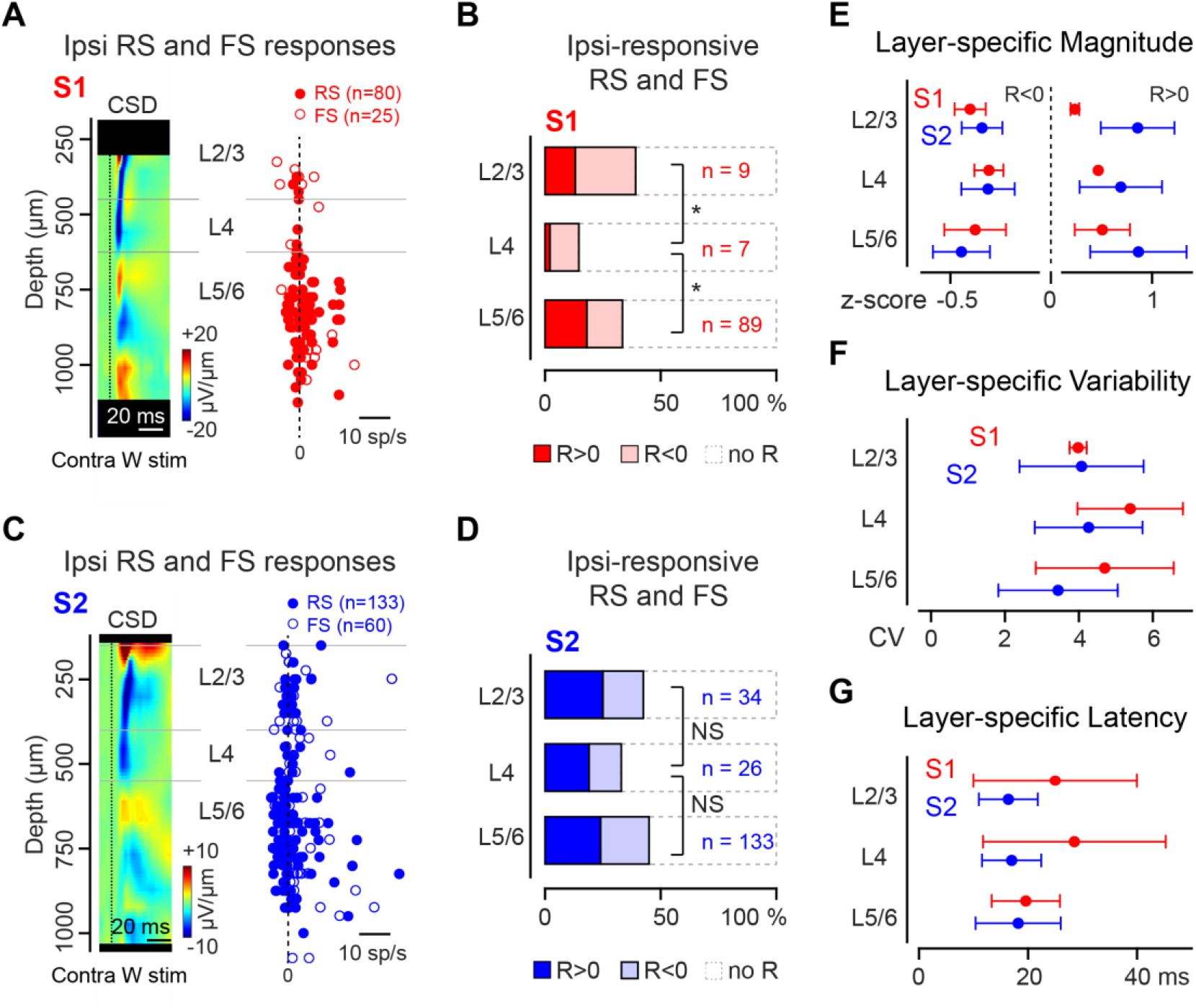
S2 ipsilateral stimulus-responsive RS and FS neurons are found in all laminae. (**A**) (Right) Mean response magnitude of 80 ipsilateral stimulus-responsive S1 RS neurons (filled markers) and 25 FS neurons as a function of cortical depth. (Left) Earliest sink (blue) in the current source density (CSD) map evoked by contralateral whisker stimuli reflects the location of layer 4 (L4). (**B**) Smaller proportion of stimulus-responsive RS and FS neurons in L4 of S1 compared to layer 2/3 (L2/3) and layer 5/6 (L5/6) (L2/3: 39% (9/23), L4: 15% (7/48), L5/6: 33% (89/266), L2/3 vs L4: p = 0.041, L5/6 vs L4: p = 0.018, chi-squared test with Bonferroni correction for two comparisons). (**C**) Same as (A), but for 133 ipsilateral stimulus-responsive S2 RS neurons and 60 FS neurons. (**D**) Comparable proportions of stimulus-responsive RS and FS neurons in L4 as in L2/3 and L5/6 in S2 (L2/3: 43% (34/80), L4: 33% (26/79), L5/6: 45% (133/295), L4 vs L2/3: p = 0.42, L5/6 vs L4: p = 0.10, chi-squared test with Bonferroni correction for two comparisons). (**E**) Comparable positive (R>0) and negative (R<0) ipsilateral response magnitudes across laminae in S1 and S2 (S1 R>0: L2/3: 0.24 ± 0.04 (n=3), z-score, median ± MAD, L4: 0.47 ± 0.00 (n=1), L5/6: 0.51 ± 0.27 (n=48), p = 0.69, Kruskal-Wallis test, S1 R<0: L2/3: -0.41 ± 0.08 (n=6), L4: -0.31 ± 0.07 (n=6), L5/6: -0.38 ± 0.16 (n=41), p = 0.60 / S2 R>0: L2/3: 0.87 ± 0.37 (n=20), L4: 0.70 ± 0.41 (n=15), L5/6: 0.88 ± 0.48 (n=71), p = 0.72, S2 R<0: L2/3: -0.35 ± 0.10 (n=14), L4: -0.32 ± 0.14 (n=11), L5/6: -0.45 ± 0.14 (n=62), p = 0.038, further pairwise comparisons using Tukey’s test all p>0.05). (F) Comparable ipsilateral positive and negative response variability across laminae in S1 and S2. Variability is quantified as the coefficient of variation (CV) of the response magnitude. (S1: L2/3: 3.97 ± 0.23 (n=9), median ± MAD, L4: 5.37 ± 1.42 (n=7), L5/6: 4.69 ± 1.86 (n=89), p = 0.80, Kruskal-Wallis test, S2: L2/3: 4.22 ± 1.67 (n=34), L4: 4.41 ± 1.45 (n=26), L5/6: 3.59 ± 1.61 (n=133), p = 0.48). (**G**) Comparable ipsilateral positive and negative response onset latency across laminae in S1 and S2. (S1: L2/3: 25.0 ± 15.0 ms (n=7), median ± MAD, L4: 28.5 ± 16.8 ms (n=6), L5/6: 19.6 ± 6.3 ms (n=63), p = 0.87, Kruskal-Wallis test, S2: L2/3: 16.4 ± 5.4 ms (n=21), L4: 17.0 ± 5.4 ms (n=20), L5/6: 18.2 ± 7.8 (n=106), p = 0.71). NS p≥0.05, * p<0.05.

### S2 neuron spiking supports higher ipsilateral stimulus decoding accuracy

Given that ipsilateral whisker deflections elicit relatively small amplitude and more variable increases and decreases in spiking in a fraction of RS neurons in S1 and S2 compared to contralateral whisker stimuli, it is unclear how accurately neuronal population activity enables single-trial ipsilateral stimulus decoding. To answer this question, we first probed whether the occurrence of an ipsilateral stimulus delivered at a velocity of 1000 °/s could be detected from the spiking activity of populations of S1 and S2 RS neurons. We implemented linear discriminant analysis (LDA) classifiers to partition RS neuron spike counts occurring 50 ms post stimulus, and compared the same intervals on trials with no stimulus (Figure 5A, see Materials and Methods). It is important to note that LDA allows individual neurons to contribute to stimulus detection independently and regardless of the sign of their stimulus-evoked spiking modulation. This means that both positive and negative changes in spiking may contribute to stimulus detection assuming they provide useful information to the classifier. Simultaneously recorded RS neurons from 8 S1 and 8 S2 recordings were initially used as input to the classifiers (S1: 22 ± 4 RS/rec, median ± MAD, range: 14 – 35 RS/rec, S2: 27 ± 9 RS/rec, range: 14 – 40 RS/rec, p = 0.55, two-sided Wilcoxon rank-sum test), which we refer to here as *within-recording* classifiers. For each neuron, stimulus-evoked and spontaneous spike counts from 70 trials each were randomly assigned to a training or a testing set according to a 10-fold cross-validation scheme, resulting in a total of 126 training trials and 14 testing trials. We found that the presence of an ipsilateral whisker stimulation could be detected with above-chance accuracy using either S1 or S2 RS neuron spiking (S1: 56.4 ± 5.0 %, chance: 49.6 ± 1.8 %, median ± MAD, p = 0.039, two-sided Wilcoxon rank-sum test, S2: 72.9 ± 11.1 %, chance: 51.1 ± 1.8 %, p = 0.0078), but with higher performance from S2 than S1 populations (p = 0.041, two-sided Wilcoxon rank-sum test) (Figure 5B).

**Figure 5.**
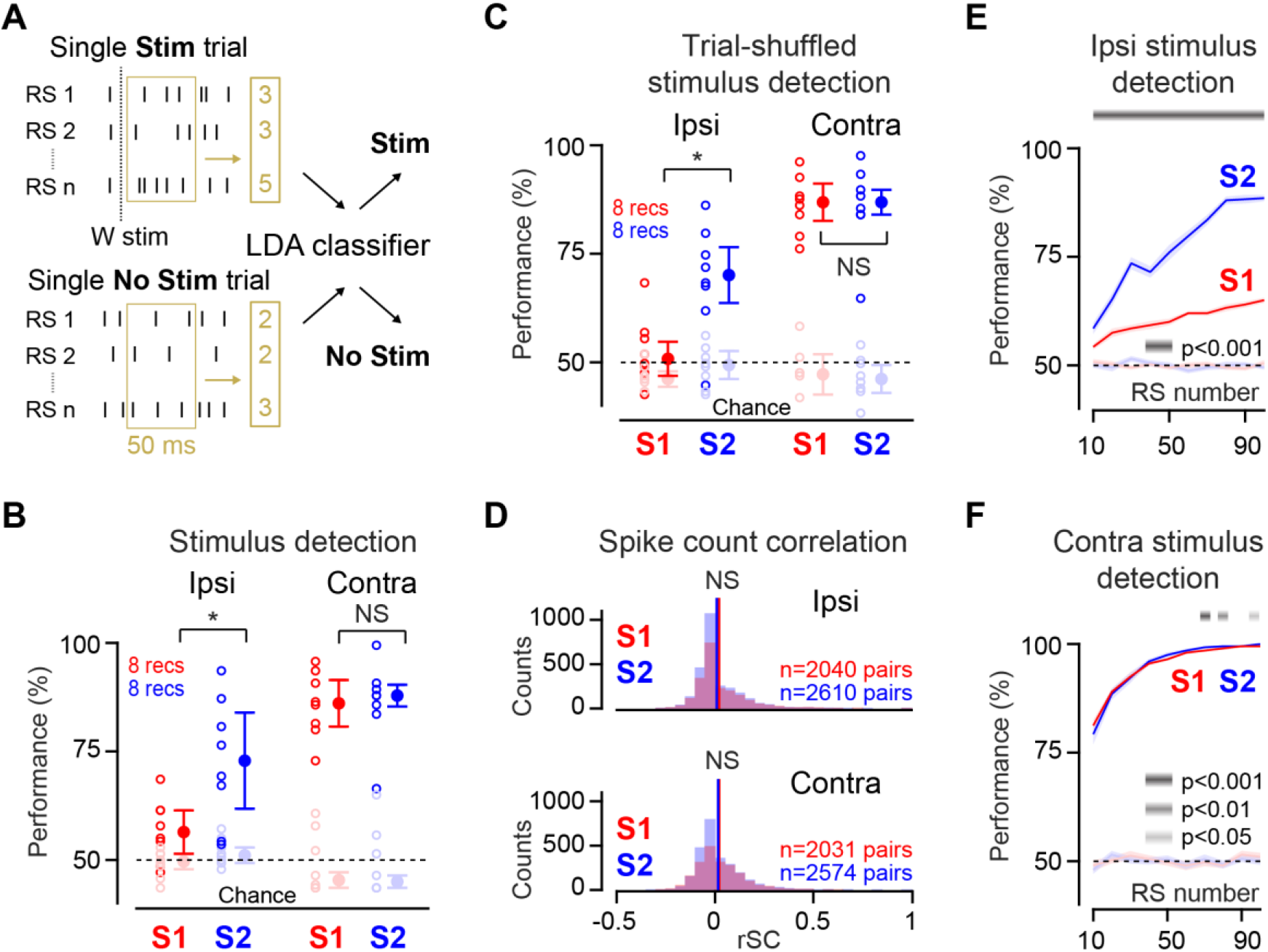
Higher ipsilateral stimulus detectability from S2 RS neuron spiking compared to S1. (A) A linear discriminant analysis (LDA) classifier partitions RS neuron single-trial spike counts occurring between 0 and 50 ms after whisker stimulation (Stim) from spike counts measured in the absence of whisker stimuli (No Stim). (**B**) Higher ipsilateral stimulus detection performance for simultaneously recorded S2 RS neurons compared to S1 RS neurons and comparable detection performance for contralateral stimuli (Ipsi: S1: 56.4 ± 5.0 % (n= 8 recordings), S2: 72.9 ± 11.1 % (n = 8 recordings), median ± MAD, p = 0.041, two-sided Wilcoxon rank-sum test, Contra: S1: 86.1 ± 5.4 %, S2: 87.9 ± 2.5 %, p = 0.74). (**C**) Same as (B) but for RS neuron spike counts randomly shuffled across trials (Ipsi: S1: 53.2 ± 3.9 % (n= 8 recordings), S2: 72.5 ± 6.4 % (n = 8 recordings), median ± MAD, p = 0.021, two-sided Wilcoxon rank-sum test, Contra: S1: 89.3 ± 4.3 %, S2: 89.3 ± 2.9 %, p = 0.74). (**D**) Comparable magnitude of spike count correlation (rSC) between pairs of RS neurons in S1 and S2. Vertical bars represent the mean rSC values (Ipsi: S1: 0.020 ± 0.123, mean ± SD, S2: 0.009 ± 0.109, p = 0.21, two-sided *t*-test, Contra: S1: 0.021 ± 0.115, S2: 0.016 ± 0.111, p = 0.14). (**E**) S1 and S2 ipsilateral stimulus detection performance (median ± SD) as a function of the number of RS neurons selected across recordings as inputs to the classifier. Grey bar at the top indicates significance of the S1 versus S2 comparison using a two-sided Wilcoxon rank-sum test. (**F**) Same as (E), but for contralateral stimulus detection performance. Chance datapoints are obtained by randomly shuffling the class labels of the training set trials. All detection performances are larger than performances obtained for chance data (p<0.05, two-sided Wilcoxon signed-rank test). In (E and F), median detection performance is computed across 100 repetitions of the classification task, and error bars represent a bootstrapped estimate of the standard deviation of the median. NS p≥0.05, * p<0.05.

This difference was specific to the ipsilateral stimulus, since contralateral stimuli were detected equally well from S1 or S2 spiking (S1: 86.1 ± 5.4 %, median ± MAD, S2: 87.9 ± 2.5 %, p = 0.74, two-sided Wilcoxon rank-sum test), and with higher overall accuracy (Figure 5B). To further examine the contribution of specific subpopulations of RS neurons to stimulus detection, we implemented a different set of classifiers, selecting the classifier input RS neuron population by random sampling across recordings, which we refer to here as *across-recordings* classifiers. Across-recordings sampling abolishes trial-by-trial covariations in individual neuron activity, which may affect stimulus coding (Zohary et al. 1994, reviewed in Averbeck et al. 2006). As a control, we first verified that the S1 versus S2 difference in ipsilateral stimulus detection performance was not driven by a difference in such covariations. First, we built *within-recording* classifiers as described above, except that we randomly shuffled spike counts across trials, thereby disrupting trial-specific covariations in spiking across simultaneously recorded neurons. Doing so did not eliminate the S1 versus S2 difference in ipsilateral stimulus detection performance, and preserved the comparable detection performances in S1 and S2 obtained for contralateral stimuli (Figure 5C). Then, we directly estimated the amount of covariation in the stimulus- evoked activity of individual neurons by measuring spike count correlations (rSC) across pairs of simultaneously recorded neurons, and found no difference in their magnitude comparing S1 and S2 for either ipsilateral or contralateral stimuli (Figure 5D). Having established that covariations in individual neuron activities on a trial-by-trial basis do not differentially affect S1 and S2 stimulus detection performances, we built *across- recordings* classifiers selecting RS neurons forming the classifier input population from all 8 S1 and 8 S2 recordings respectively (selection pool size of 184 RS neurons for S1 and 205 RS neurons for S2), and used a diagonal covariance matrix to prevent the contribution of spurious covariations in spiking activity to detection performance. Similarly to the classifiers built from simultaneously recorded neurons, stimulus-evoked and spontaneous spike counts were randomly assigned to a training or a testing set, according to a 10-fold cross-validation scheme. Within each set, spike counts were sampled with replacement to generate a total of 180 training trials and 20 testing trials. This procedure was repeated 100 times to determine detection performance across different combinations of neurons and trials. Such classifiers recapitulated the S1 versus S2 difference in ipsilateral stimulus detection performance, and the comparable detection performances obtained for contralateral stimuli (Figure 5E, F). These results were independent of the number of trials and repetitions, though absolute detection performance values in both S1 and S2 increased with the number of neurons forming the classifier input population for both ipsilateral and contralateral stimuli (Figure 5E, F).

To specifically examine the contribution of subpopulations of RS neurons located in different neocortical laminae to stimulus detection, we built classifiers with an input population size of 10 RS neurons to account for the size of laminar-specific S1 and S2 sampling pools (see Materials and Methods). We found that in both S1 and S2, L5/6 neurons lead to greatest ipsilateral stimulus detection performance (Figure 6A), while they performed worst for contralateral stimulus detection (Figure 6B).

**Figure 6.**
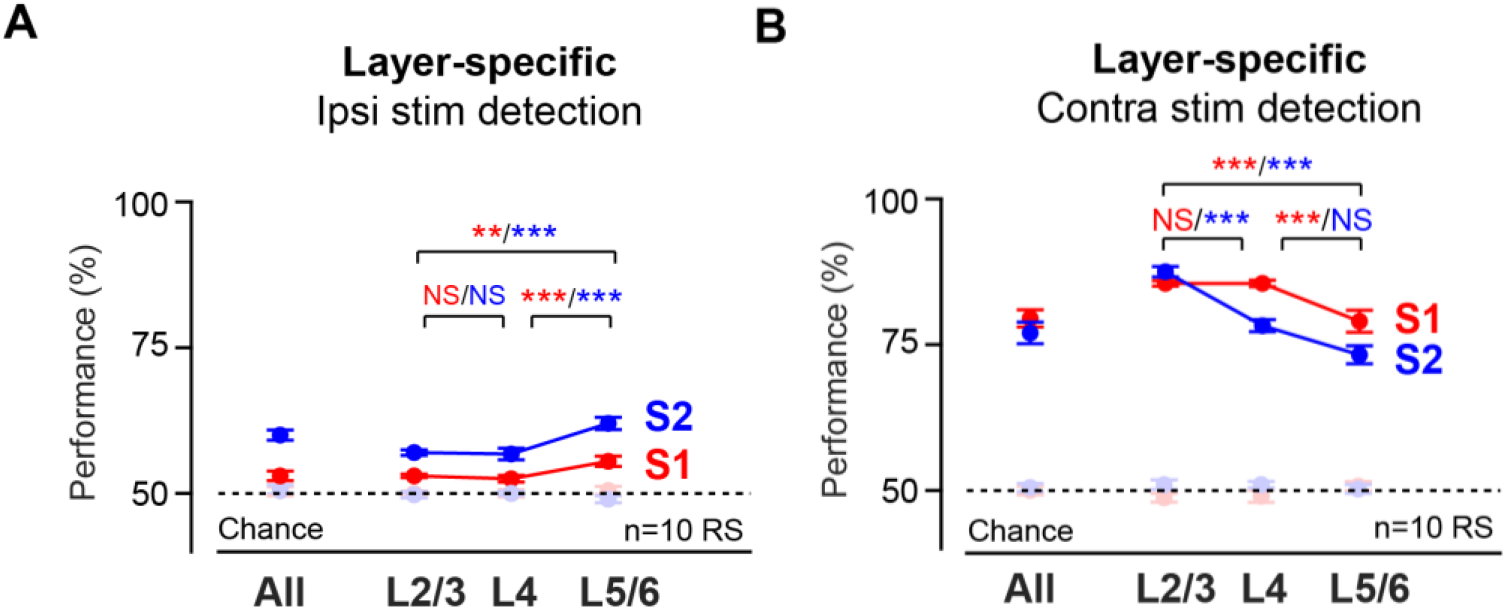
Highest ipsilateral stimulus detectability from L5/6 RS neuron. (**A**) Higher ipsilateral stimulus detection performance for L5/6 RS neurons compared to L2/3 and L4 neurons in S1 and S2 (S1: L2/3: 53 ± 0.3 %, L4: 52.5 ± 0.6 %, L5/6: 55.5 ± 0.9 %, median ± SD, L5/6 vs L2/3: p= 0.0012, L5/6 vs L4: p = 0.00016, two-sided Wilcoxon rank-sum test with Bonferroni correction for three comparisons, S2: L2/3: 57 ± 0.5 %, L4: 56.8 ± 1 %, L5/6: 62 ± 1 %, L5/6 vs L2/3: p = 1.43·10^-6^, L5/6 vs L4: p = 0.00027). (**B**) Higher contralateral stimulus detection performance for L2/3 RS neurons compared to L5/6 RS neurons in S1 and S2 (S1: L2/3: 85.5 ± 0.5 %, L5/6: 79 ± 2 %, median ± SD, p= 1.96·10^-6^, two-sided Wilcoxon rank-sum test, S2: L2/3: 87.5 ± 0.9 %, L5/6: 73.3 ± 1.5 %, p = 1.90·10^-11^). Chance datapoints are obtained by randomly shuffling the class labels of the training set trials. All detection performances are larger than performances obtained for chance data (p<0.01, two-sided Wilcoxon signed-rank test). Median detection performance is computed across 100 repetitions of the classification task, and error bars represent a bootstrapped estimate of the standard deviation of the median. NS p≥0.05, ** p<0.01, *** p<0.001.

One possible explanation for the higher detection performance obtained from S2 spiking within and across laminae could be the higher proportion of stimulus-responsive RS neurons observed in S2 compared to S1. To investigate this, we implemented classifiers with 24 input RS neurons (our average yield per recording), while matching the proportion of stimulus-responsive neurons in S1 and S2, which led to a comparable interareal difference in detection performance (Figure 7A), thus ruling out the number of stimulus-responsive RS neurons in each area as a contributor of the S2 versus S1 difference in detection performance. Then, we focused on stimulus-responsive RS neurons, as the spiking of non-responsive RS neurons led to detection performances not different from chance levels in both S1 and S2 (Figure 7B). Detection performance diminished, and more so in S2 than in S1, when only RS neurons with negative response magnitudes were used as input to the classifier (Figure 7C), which resulted in a drastic reduction of the amplitude of the S2 versus S1 difference in stimulus detectability (Figure 7D). On the contrary, when only RS neurons with positive response magnitudes were used as the classifier input population, detection performance was further enhanced in S2 compared to S1 (Figure 7C), leading to an augmentation of the interareal difference in stimulus detection performance (Figure 7D), likely due to the larger absolute magnitude of positive responses, and even more so in S2, as compared to the negative ones. This finding thus identifies a predominant role for S1 and S2 RS neurons with increased stimulus-evoked spiking in the detection of ipsilateral stimuli, as well as in the larger detection performance achieved from S2 compared to S1.

**Figure 7.**
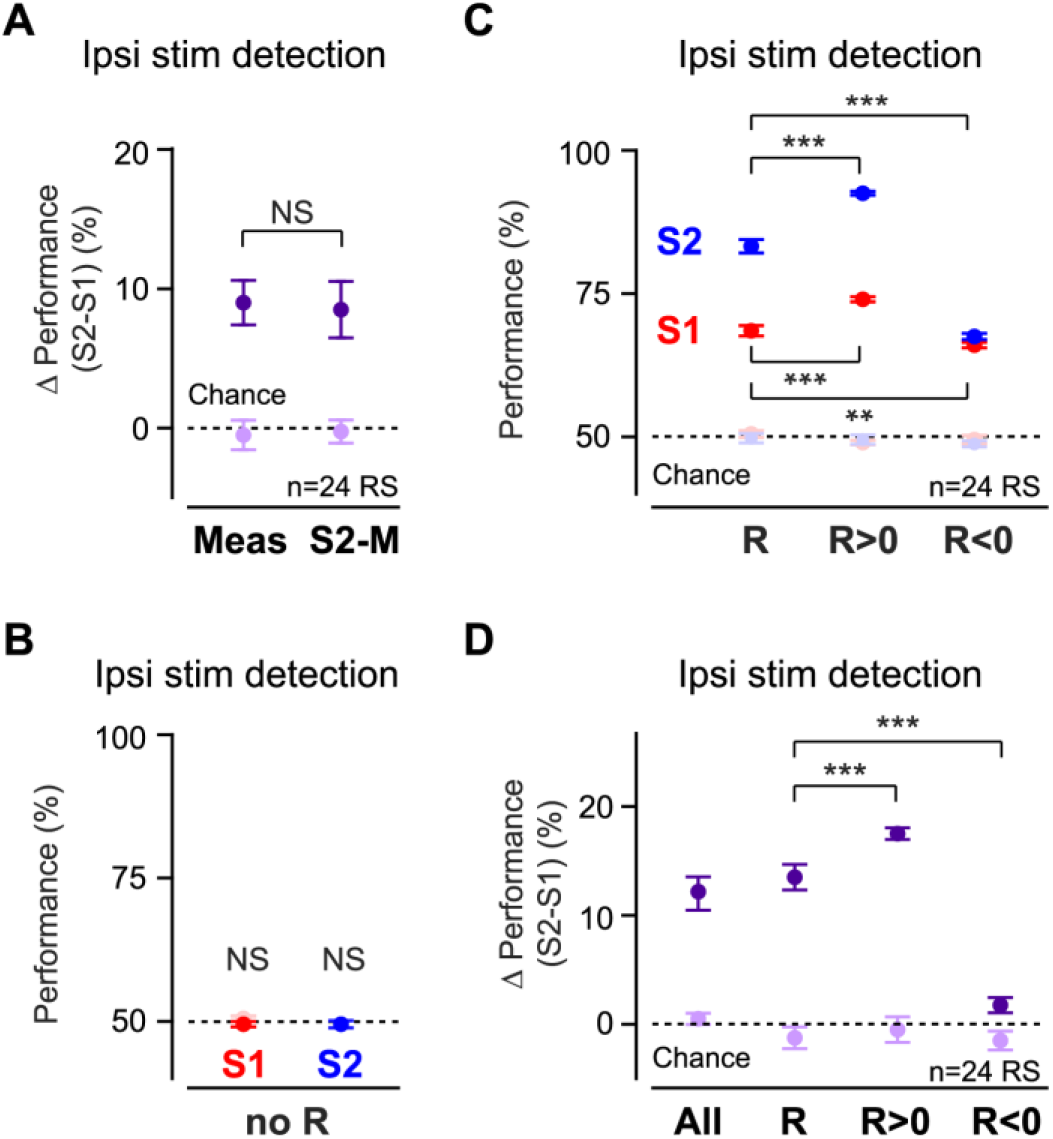
S2 versus S1 ipsilateral stimulus detectability difference arises from spiking of stimulus-responsive RS neurons with positive response magnitude. **(A)** Matching the percentage of S1 stimulus-responsive RS neurons to that measured in S2 (S2-M) does not reduce the difference in ipsilateral detection performance between the two areas (Meas) (Δ Measured %: 9 ± 1.6 %, Δ S2-Matched %: 8.5 ± 2.0 %, median ± SD, p = 0.57, two-sided Wilcoxon rank-sum test). **(B)** Chance-level ipsilateral stimulus detection performance for non-stimulus-responsive S1 and S2 RS neurons (S1: 50.0 ± 0.5 %, chance: 50.5 ± 0.4 %, median ± SD, p = 0.37, two-sided Wilcoxon rank-sum test, S2: 49.5 ± 0.6 %, chance: 49.5 ± 0.6 %, p = 0.90). **(C)** Enhanced ipsilateral stimulus detection performance for stimulus-responsive RS neurons with positive response magnitude (R>0) and decreased performance for RS neurons with negative response magnitude (R<0) (S1: R: 68.5 ± 0.9 %, R>0: 74 ± 0.4 %, R<0: 66 ± 0.5 %, median ± SD, R versus R>0: p = 2.78·10^-10^, R versus R<0: p = 0.0044, two-sided Wilcoxon rank-sum test with Bonferroni correction for two comparisons, S2: R: 83.3 ± 1.1 %, R>0: 92.5 ± 0.4 %, R<0: 67.5 ± 0.5 %, R versus R>0: p = 4.03·10^- 16^, R versus R<0: p = 1.12·10^-26^). **(D)** Increased S2 versus S1 difference in ipsilateral stimulus detection performance for stimulus-responsive RS neurons with positive response magnitude (R>0) and strong reduction for stimulus-responsive RS neurons with negative response magnitude (R<0) (Δ R: 13.5 ± 1.1 %, Δ R>0: 17.5 ± 0.6 %, Δ R<0: 1.8 ± 0.7 %, median ± SD, Δ R vs Δ R>0: p = 0.00064, Δ R vs Δ R<0: p = 1.58·10^-16^, two-sided Wilcoxon rank-sum test with Bonferroni correction for two comparisons). Median detection performance is computed across 100 repetitions of the classification task. Error bars represent a bootstrapped estimate of the standard deviation of the median. Chance datapoints are obtained by randomly shuffling the class labels of the training set trials. All detection performances, and S2-S1 detection performance differences, are larger than performances obtained for chance data (p<0.01, two-sided Wilcoxon rank-sum test). NS p≥0.05, ** p<0.01, *** p<0.001.

Having established that the spiking activity of both S1 and S2 RS neurons contains enough information to detect the occurrence of single ipsilateral stimuli, we then probed whether it contained additional information regarding stimulus features. We focused on whisker deflection velocity, as this has previously been shown to be an important aspect of contralateral whisker motion that is encoded in the spiking rate of individual S1 neurons (Simons 1978, Ito 1985, Pinto et al. 2000, Arabzadeh et al. 2003, Arabzadeh et al. 2004, Wilent and Contreras 2004, Boloori et al. 2010, Ranjbar-Slamloo and Arabzadeh 2017).

We asked whether single whisker stimuli delivered at two different velocities could be discriminated on the basis of ipsilateral neural activity in S1 versus S2. We considered changes in RS neurons spiking evoked by stimuli of 200 °/s and 1000 °/s velocity (Figure 8A). The 200 °/s ipsilateral stimuli elicited a significant change in firing in a proportion of S1 and S2 RS neurons (Table 2) comparable to that obtained in response to 1000 °/s stimuli (Table 1) (S1: p = 0.55, S2: p = 0.60, chi-squared test). Overall, RS responses to 200 °/s ipsilateral stimuli were characterized by a reduction in magnitude, without noticeable change in variability or onset latency compared to responses evoked by 1000 °/s stimuli (Tables 1 and 2). Exceptions included the magnitude of positive responses in S1 RS neurons which was similar for the two stimulus velocities (p = 0.34, one-sided Wilcoxon rank-sum test, all other comparisons: p < 0.05), and the variability of negative responses in S2 RS neurons which was larger for 200 °/s stimuli than for 1000 °/s stimuli (p = 0.0027, one-sided Wilcoxon rank-sum test, all other comparisons: p ≥ 0.05). First, we investigated whether such overall smaller evoked changes in activity could still support ipsilateral stimulus detection. As we previously showed that implementing classifiers from neurons sampled across recordings did not noticeably alter decoding performances, we again built *across-recordings* classifiers by randomly selecting 24 RS neurons across 6 S1 and 6 S2 recordings (selection pool size of 169 S1 neurons and 144 S2 neurons) and found that the presence of weaker 200 °/s ipsilateral stimuli could indeed be detected from the spiking activity of S1 and S2 RS neurons (Figure 8B) (S1: 53.8 ± 0.8 %, chance: 50.5 ± 0.4 %, median ± SD, p = 2.48·10^-5^, two-sided Wilcoxon rank-sum test, S2: 61.5 ± 1.3%, chance: 49.5 ± 0.7 %, p = 1.43·10^-24^). Then, we trained and cross-validated classifiers with 24 input RS neurons selected from a pool of 83 S1 RS neurons or 77 S2 RS neurons obtained from 3 S1 and 3 S2 recordings respectively, during which stimuli of both velocities were delivered (Figure 8C).

**Figure 8.**
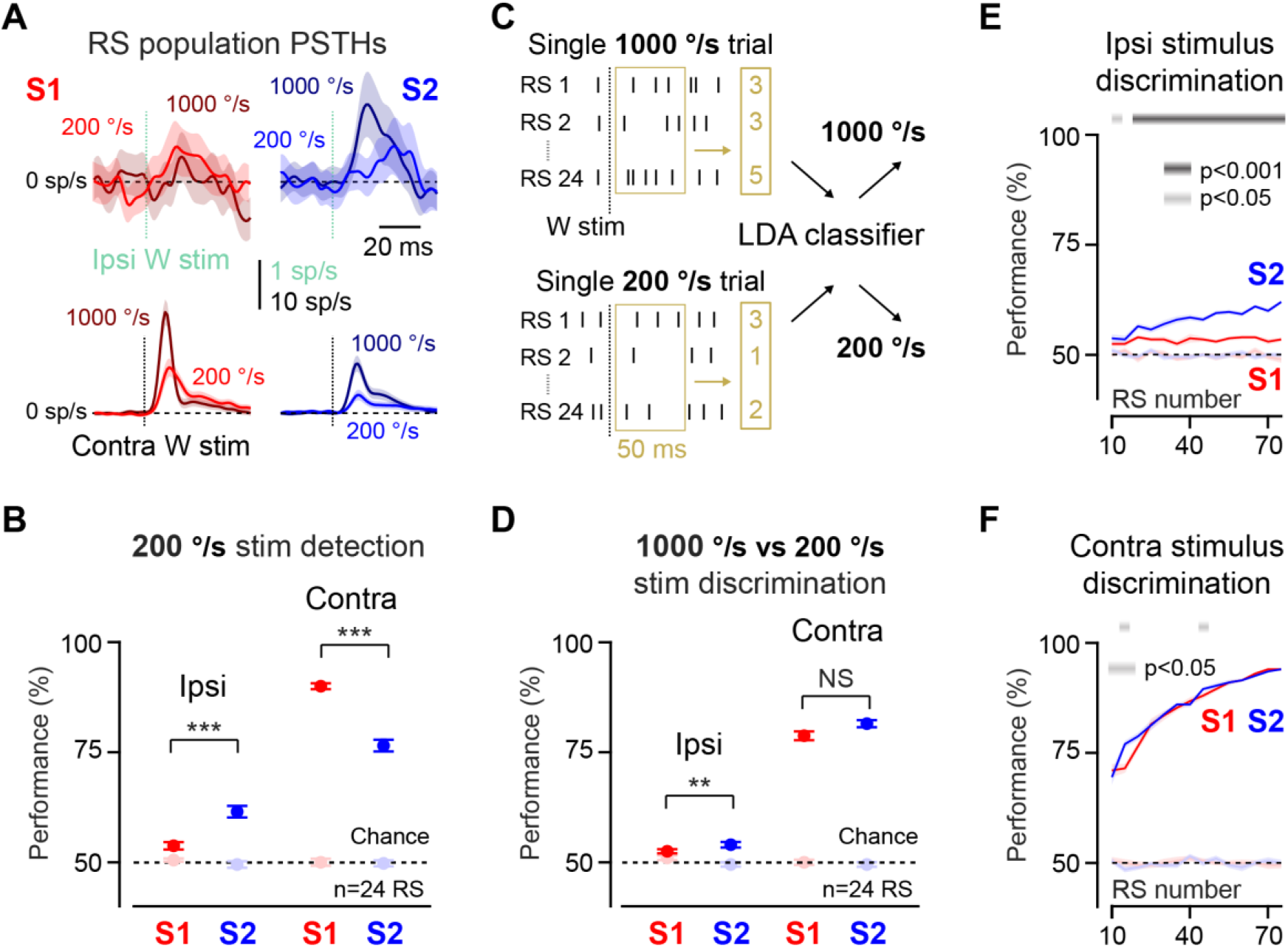
Higher ipsilateral stimulus discriminability from S2 RS neuron spiking compared to S1. (A) (Top) Baseline-subtracted population PSTHs for S1 (red) and S2 (blue) RS neurons in response to 200 °/s (n = 172 S1 RS, n=148 S2 RS) and 1000 °/s (n = 263 S1 RS, n=359 S2 RS) ipsilateral stimuli. (Bottom) Same for contralateral stimuli. Shaded areas represent the SEM. (**B**) 200 °/s ipsilateral stimuli are detectable from S1 and S2 spiking at above-chance performance. Detection performance is higher for S2 than for S1 and lower than for contralateral stimuli (Ipsi: S1: 53.8 ± 0.8 %, S2: 61.5 ± 1.3 %, median ± SD, p = 3.75·10^-10^, two-sided Wilcoxon rank-sum test, Contra: S1: 90 ± 0.7 %, S2: 76.5 ± 1.3 %, p = 7.04·10^-24^, Ipsi vs Contra: S1: p = 2.92·10^-34^, S2: p = 1.19·10^-21^). (**C**) A linear discriminant analysis (LDA) classifier partitions single-trial spike counts from 24 randomly chosen RS neurons, measured between 0 and 50 ms after 1000 °/s and 200 °/s whisker stimuli. (**D**) Above-chance discrimination performance for ipsilateral stimuli of different velocities from S1 and S2 RS neuron spiking. Higher discriminability in S2 compared to S1, though lower than contralateral stimulus discriminability (Ipsi: S1: 52.5 ± 0.5 %, S2: 54 ± 0.6 %, median ± SD, p = 0.0085, two-sided Wilcoxon rank-sum test, Contra: S1: 78.8 ± 1 %, S2: 81.5 ± 0.8 %, p = 0.087, Ipsi vs Contra: S1: p = 2.81·10^-34^, S2: p = 3.7·10^-34^). (**E**) S1 and S2 ipsilateral stimulus discrimination performance (median ± SD) as a function of the number of RS neurons used as inputs to the classifier. Grey bar at the top indicates significance of the S1 versus S2 comparison using a two-sided Wilcoxon rank-sum test. (**F**) Same as (E), but for contralateral stimulus discrimination performance. Median detection performance is computed across 100 repetitions of the classification task. Error bars represent a bootstrapped estimate of the standard deviation of the median. Chance datapoints are obtained by randomly shuffling the class labels of the training set trials. All detection performances are larger than performances obtained for chance data (p<0.01, two-sided Wilcoxon rank-sum test). NS p≥0.05, ** p<0.01, *** p<0.001.

**Table 2:**
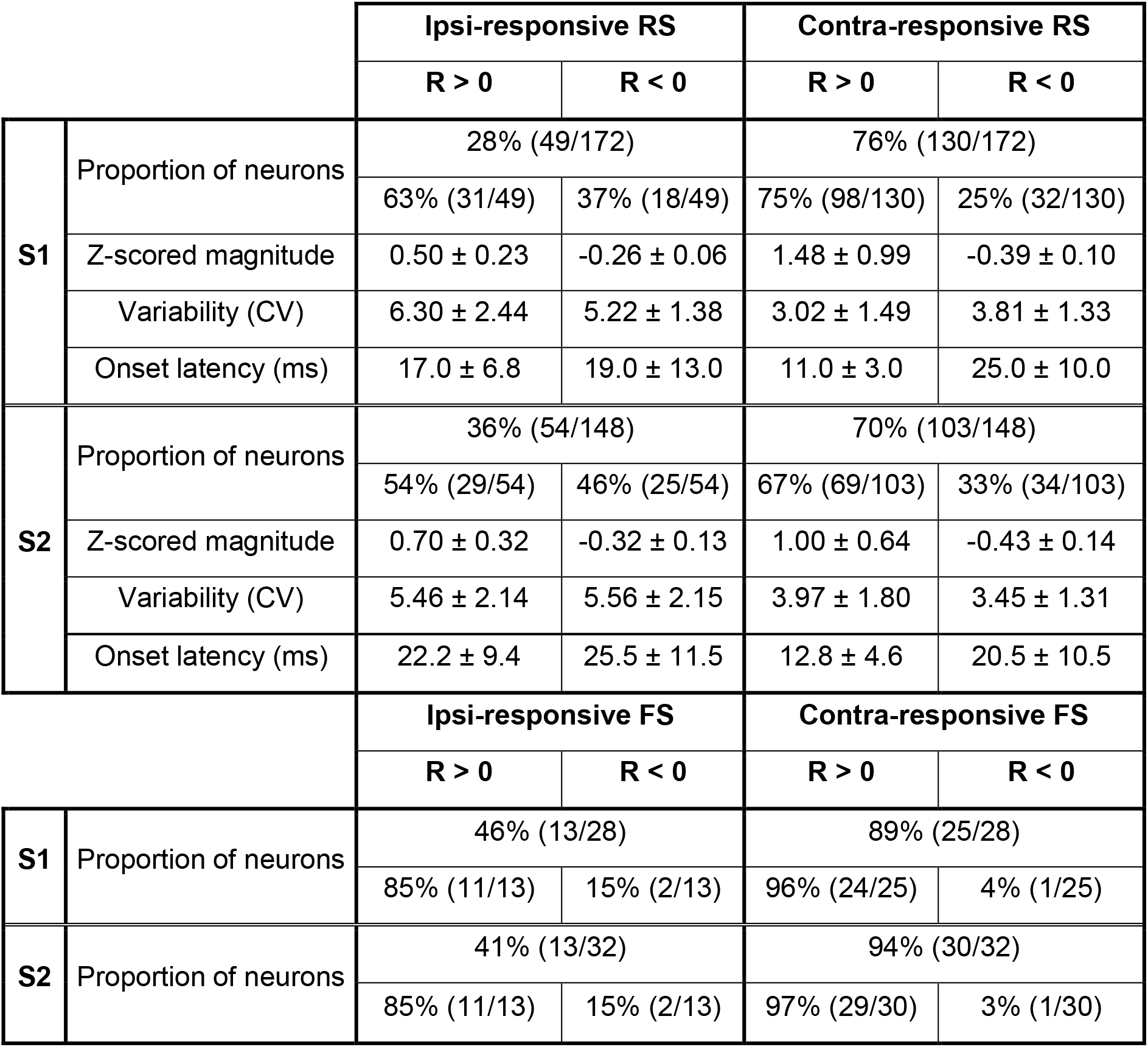
Proportion of stimulus-responsive RS and FS neurons and response properties: 200 °/s stimuli. Proportion of RS and FS neurons with a significant response to 200 °/s ipsilateral or contralateral stimuli. Response magnitude (z-score), variability (coefficient of variation, CV), and onset latency are reported as median ± MAD.

|  |  | **Ipsi-responsive RS** |  | **Contra-responsive RS** |  |
| --- | --- | --- | --- | --- | --- |
|  |  | **R > 0** | **R < 0** | **R > 0** | **R < 0** |
| **S1** | Proportion of neurons | 28% (49/172) |  | 76% (130/172) |  |
|  |  | 63% (31/49) | 37% (18/49) | 75% (98/130) | 25% (32/130) |
|  | Z-scored magnitude | 0.50 ± 0.23 | -0.26 ± 0.06 | 1.48 ± 0.99 | -0.39 ± 0.10 |
|  | Variability (CV) | 6.30 ± 2.44 | 5.22 ± 1.38 | 3.02 ± 1.49 | 3.81 ± 1.33 |
|  | Onset latency (ms) | 17.0 ± 6.8 | 19.0 ± 13.0 | 11.0 ± 3.0 | 25.0 ± 10.0 |
| **S2** | Proportion of neurons | 36% (54/148) |  | 70% (103/148) |  |
|  |  | 54% (29/54) | 46% (25/54) | 67% (69/103) | 33% (34/103) |
|  | Z-scored magnitude | 0.70 ± 0.32 | -0.32 ± 0.13 | 1.00 ± 0.64 | -0.43 ± 0.14 |
|  | Variability (CV) | 5.46 ± 2.14 | 5.56 ± 2.15 | 3.97 ± 1.80 | 3.45 ± 1.31 |
|  | Onset latency (ms) | 22.2 ± 9.4 | 25.5 ± 11.5 | 12.8 ± 4.6 | 20.5 ± 10.5 |
|  |  | **Ipsi-responsive FS** |  | **Contra-responsive FS** |  |
|  |  | **R > 0** | **R < 0** | **R > 0** | **R < 0** |
| **S1** | Proportion of neurons | 46% (13/28) |  | 89% (25/28) |  |
|  |  | 85% (11/13) | 15% (2/13) | 96% (24/25) | 4% (1/25) |
| **S2** | Proportion of neurons | 41% (13/32) |  | 94% (30/32) |  |
|  |  | 85% (11/13) | 15% (2/13) | 97% (29/30) | 3% (1/30) |

Overall, S1 and S2 velocity discrimination performances were lower than values obtained for contralateral stimuli (Figure 8D), but significantly higher than chance (S1: 52.5 ± 0.5 %, chance: 51 ± 0.5 %, median ± SD, p = 0.0027, two-sided Wilcoxon rank-sum test, S2: 54 ± 0.6 %, chance: 49.5 ± 0.5 %, p = 3.75·10^-9^). In the same way that S2 spiking supported higher ipsilateral stimulus detectability, it also enabled higher ipsilateral stimulus velocity discriminability compared to S1 (S1 versus S2: p = 0.0085, two-sided Wilcoxon rank-sum test). These results were independent of the number of trials chosen to train and test the classifiers, and of the number of repetitions of the classification task. Increasing the number of RS neurons forming the classifier input population led to an increase in ipsilateral stimulus discrimination performance in S2 only, therefore further enhancing the S2 versus S1 difference in ipsilateral stimulus discriminability (Figure 8E), while both S1 and S2 contralateral stimulus discrimination performances increased as a function of the size of the classifier input population (Figure 8F). Together, our classifier-based analyses support a role for the activity of RS neurons in somatosensory cortices, in particular in S2, in encoding the presence and the velocity of ipsilateral tactile stimuli in addition to representing contralateral sensory information.

## Discussion

Our results revealed a strong representation of ipsilateral tactile stimuli in S1 and S2 of the awake mouse brain. Although spikes from both S1 and S2 RS neurons enabled the decoding of ipsilateral tactile stimuli, S2 spikes led to greater stimulus detection and feature discrimination. Ipsilateral stimuli elicited increases and decreases in spiking with equal probability in S1 and S2, both contributing to stimulus decoding, yet higher performance in S2 could be explained by less variable and larger stimulus-evoked increases in spike rate compared to S1. In S1 and S2, such ipsilateral encoding of tactile information was distributed across 30-40% of neurons, located in all neocortical laminae in S2, but tending to spare layer 4 in S1. Our findings provide a functional role for ipsilateral activity in contributing to the encoding of tactile information arising from one side of the body across both cerebral hemispheres.

Our measurements conducted in the whisker system of awake mice corroborate previous findings in anesthetized rodents, and provide new insights into the cellular substrates of ipsilateral responses. Earlier studies have reported the existence in S1 and S2 of the ipsilateral hemisphere of excitatory neurons with increased spiking in response to unilateral tactile stimulation of various body parts (Carvell and Simons 1986, Armstrong-James and George 1988, Shuler et al. 2001, Genna et al. 2018). Here, we showed that during wakefulness, ipsilateral tactile stimuli elicited both increased and decreased spiking with equal probability in RS neurons of S1 and S2. These findings contrast with sensory responses we and others measured in response to contralateral stimuli in these two regions, which occurred with higher probability, faster latency, and with a principally increased firing rate (Crochet and Petersen 2006, Yamashita et al. 2013, Minamisawa et al. 2018, Ranjbar-Slamloo and Arabzadeh 2019).

In addition, we provided a detailed characterization of ipsilateral responses in FS neurons, which are the most common subtype of neocortical GABAergic neurons in the mouse (Rudy et al. 2011). We found that FS neurons tended to be more responsive to ipsilateral stimuli than RS neurons, especially in S2, and that they overwhelmingly responded through an increase in spiking, with a faster onset latency compared to that of RS neurons. Although FS neurons may potentially mediate the decrease in spiking in RS neurons, further experimental investigations are necessary to establish a causal role. As the callosal projections thought to propagate changes in neural activity across hemispheres have been shown to be largely glutamatergic (see review by Conti and Manzoni 1994), the suppression of ipsilateral RS neuron spiking is unlikely to occur through their direct monosynaptic effect. Besides FS neurons, other subtypes of GABAergic neurons could provide feedforward inhibition. For instance, GABAergic neurons of layer 1, which express the ionotropic serotonin receptor 5HT3a (Lee et al. 2010), have been found to receive callosal inputs in the hindlimb region of S1 (Palmer et al. 2012).

Using laminar silicon probes, we were able to compare responses to ipsilateral stimuli across neocortical layers. We found that stimulus-responsive neurons were evenly located across all layers of S1 and S2, with the exception of L4 in S1, which contained a reduced number of neurons mostly exhibiting negative sensory responses. In rodents, as callosal axon terminals are known to be sparse in L4 of S1 (Wise 1975, Wise and Jones 1976, Akers and Killackey 1978), and as thalamocortical inputs targeting L4 have been shown to relay sensory signals solely from the opposite side of the body (Smith 1973, Waite 1973, Erzurumlu and Killackey 1980, Castejon et al. 2021), these rare L4 negative responses are likely mediated by translaminar feedforward inhibition. Further quantifications of ipsilateral sensory response properties did not reveal differences in response magnitude, variability, or onset latency as a function of laminar location, neither in S1, nor in S2. These results are at odds with in-vitro findings reporting larger monosynaptic excitatory postsynaptic potentials in L5 compared to L2/3 pyramidal neurons of S1 in response to callosal input activation (Petreanu et al. 2007), and with our decoding analyses that revealed a higher ipsilateral stimulus detection performance from L5/6 RS neurons compared to L2/3 and L4 neurons. This discrepancy could potentially be explained by the relatively small number of recorded stimulus-responsive neurons, especially in L2/3 and L4 of S1, affecting the robustness of comparisons across layers.

Implementing classifier-based analyses enabled us to quantify the role of relatively sparse bidirectional ipsilateral changes in spiking for stimulus coding. Our population-based decoding approach clearly revealed that both strong and weak ipsilateral whisker deflections could be detected from S1 and S2 activity, and that both increased and decreased spiking contributed to stimulus detectability. Detection performance for ipsilateral stimuli in S2 reached 85-90% for >80 input neurons, but remained moderate in S1, plateauing at around 60-65% for strong stimuli when 60 or more neurons were used as input to the classifier. These differences reflect both the magnitude of responses as well as the variability in each region. As a reference, ipsilateral stimuli were always detected with lower accuracy than contralateral ones, which, when delivered at lower velocity, were better detected from S1 than from S2 (Kwon et al. 2016). Stimulus discrimination proved less accurate than stimulus detection, though still reaching above-chance levels, suggesting that some amount of information about stimulus velocity is nonetheless contained in ipsilateral spiking activity, and in particular in the magnitude of the stimulus-evoked changes in spiking. Similar to what was found for stimulus detection, ipsilateral stimulus discrimination performance never reached that obtained for contralateral stimuli. This matches results obtained in single S1 neurons in anesthetized rats showing that 300-ms long spatiotemporal stimulation patterns are discriminated with lower accuracy when applied to a single digit of the ipsilateral forepaw than when applied to a contralateral digit (Genna et al. 2018).

A key finding of our work is the more robust representation of ipsilateral stimuli paralleled by the higher decoding performances obtained in S2 compared to S1. As we did not perform simultaneous recordings in S1 and S2, but in some instances nonetheless sequentially recorded from S1 and S2 in the same animal, we cannot completely rule out any recording- and animal-specific effects. Yet, our results align with findings in macaque monkeys showing that the proportion of neurons responding to stimulation of the ipsilateral hand increases across subsequent neocortical areas of the somatosensory pathway, with area 3b of S1 containing almost no stimulus-responsive neurons, area 2 of S1 displaying a small fraction of stimulus-responsive neurons, and S2 exhibiting a majority of stimulus-responsive neurons (Iwamura et al. 1994, Burton et al. 1998, Iwamura et al. 2001, Taoka et al. 2016). Such findings are typically explained in the light of the anatomical organization of callosal projections, with dense callosal projections between the hand regions of S2 across hemispheres and sparser projections between the equivalent regions in areas 2 and 3b of S1 (Killackey et al. 1983, Manzoni et al. 1984, Manzoni et al. 1986, Krubitzer and Kaas 1990). In the mouse, although callosal projections are known to exist between the whisker regions of both S1 and S2 across hemispheres (White and DeAmicis 1977, Carvell and Simons 1987, Olavarria and Van Sluyters 1995, Aronoff et al. 2010, Oh et al. 2014), it is unclear whether they follow a comparable organization. Separately, it is important to note that changes in activity induced by ipsilateral stimuli may be mediated by polysynaptic pathways within the recorded hemisphere, either across laminae within S1 or S2, but also across S1 and S2 as was shown for contralateral stimuli (Minamisawa et al. 2018), a topic for further investigations. Functionally, S2 neuron spikes have been shown to encode the frequency of vibrotactile stimuli, as well as object textures and shapes (Romo et al. 2002, Zuo et al. 2015, reviewed in Delhaye et al. 2018), as a result of the integration of more basic stimulus features across time and space within a given body side (Goldin et al. 2018). Our results further suggest that the spatial integration of stimulus features in S2 may go beyond contralateral inputs and encompass both body sides.

Taken together, our results reveal a cellular, laminar, and hierarchical specialization for ipsilateral tactile stimulus encoding in mouse S2 versus S1. While this makes possible the notion of a sensory representation that is distributed across the two hemispheres, callosal transection studies (Stamm and Sperry 1957, Ebner and Myers 1962, reviewed in Glickstein and Berlucchi 2008) seem to rather suggest that the ipsilateral representation is redundant, perhaps providing a substrate for the rapid transfer of learned unilateral tactile behaviors across body sides. The relevance of ipsilateral activity in S1, S2, and beyond, is likely to be further understood by investigating bilateral somatosensation, as ipsilateral tactile signals have been shown to modulate contralateral sensory responses already in S1 (Burton et al. 1998, Shuler et al. 2001, Wiest et al, 2005, Reed et al. 2011, reviewed in Tame et al. 2016). Future studies must thus be designed to probe and manipulate neocortical somatosensory activity during bilateral behavioral paradigms that engage, and rely on, the integration of contralateral and ipsilateral tactile information.

## Acknowledgements

This work was supported by the Swiss National Science Foundation postdoctoral fellowships P2ELP3_168506 and P300PA_177861 (AP), the National Institute of Neurological Disorders and Stroke BRAIN Grant R01NS104928 (GBS), and the National Institute of Neurological Disorders and Stroke Grant R21NS112783 (AP and GBS). We thank Bilal Haider for sharing the Scnn1a-Tg3-Cre x LSL-ChR2 mice and for insightful feedback on the manuscript, Audrey Sederberg for valuable suggestions about the classifier-based analyses, and other members of the Stanley laboratory for helpful discussions.

